# Differential requirements of cyclase associated protein (CAP) for actin turnover during the lytic cycle of *Toxoplasma gondii*

**DOI:** 10.1101/569368

**Authors:** Alex Hunt, Jeanette Wagener, Robyn Kent, Romain Carmeille, Matt Russell, Christopher Peddie, Lucy Collinson, Aoife Heaslip, Gary E. Ward, Moritz Treeck

## Abstract

*Toxoplasma gondii* contains a limited subset of actin binding proteins. Here we show that ablation of the putative actin regulator cyclase-associated protein (TgCAP) leads to significant defects in some but not all actin dependent processes, including a defect in cell-cell communication, but surprisingly not synchronicity of division. Two CAP isoforms originate from alternative translational start sites and are beneficial for parasite fitness while a single isoform is sufficient for virulence in mice. Examination of the mutant parasites by 3D electron microscopy reveals that loss of CAP results in a defect to form a normal residual body, but all parasites remain connected within the vacuole. This dissociates synchronicity of division and parasite rosetting and reveals that establishment and maintenance of the residual body may be more complex than previously thought. These results highlight the different spatial requirements for actin turnover in *Toxoplasma*, controlled by a reduced subset of actin binding proteins.

## Introduction

*Toxoplasma gondii* is an obligate intracellular parasite, belonging to the Apicomplexa phylum. The Apicomplexa, which also include *Plasmodium* and *Cryptosporidium* species, pose a significant global public health burden. *Toxoplasma,* specifically, is one of the most prevalent human pathogens, chronically infecting ~30% of the world’s population (Swapna & Parkinson, 2017). While most infections are asymptomatic, in congenitally infected and immunocompromised patients, disease outcomes are often severe and potentially fatal (Halonen & Weiss, 2013). During acute infection of the host, the asexual tachyzoite stage of *Toxoplasma gondii* undergoes cycles of active invasion, replication and egress from host cells. This lytic cycle leads to rapid proliferation and dissemination of the parasite throughout the host (Black & Boothroyd, 2000). To facilitate these processes, *Toxoplasma* utilises a unique form of locomotion, called gliding motility, which relies on actin and an unconventional myosin motor (Frénal, Dubremetz, et al., 2017). This motor allows the parasite to actively invade host cells, where it forms a protective parasitophorous vacuole (PV). PV structural integrity and stability is sustained through the parasite’s release of dense granule proteins from secretory vesicles (Heaslip et al., 2016). Additionally, several dense granule proteins are transported into the host cell where they co-opt or interfere with host cell functions (Hakimi et al., 2017). Within the PV, *Toxoplasma* begins a unique form of cell division called endodyogeny (Sheffield & Melton, 1968). Here, two daughter cells are synchronously assembled within the mother cell before daughter cell budding (Delbac et al., 2001). This initiates at the apical pole of the mother cell and once complete, the daughter cells bud from the mother cell but remain attached at their basal pole to the residual body: a membranous compartment containing maternal remnants of organelles and cytoskeletal elements (Muñiz-Hernández et al., 2011). During cell division, organelles such as mitochondria and the apicoplast, a unique and essential organelle which contributes to isoprenoid synthesis (Vaishnava & Striepen, 2006), also divide and are distributed between the two daughter parasites. Following multiple rounds of cell division, parasites organise into a rosette-like pattern around the residual body. Formation of the characteristic rosette pattern requires actin, several myosins, and other actin-binding proteins (Frénal, Jacot, et al., 2017; Haase et al., 2015; Jacot et al., 2013; Periz et al., 2017; Tosetti et al., 2019). It has been hypothesised that the inter-parasite connections via the residual body are not only important for parasite organisation, but also play a key role in cell-cell communication. Such communication was measured by the transfer of reporter proteins between parasites and is believed to ensure the synchronous division of parasites within a vacuole (Frénal, Jacot, et al., 2017). The parasites continue to replicate until host cell lysis and parasite egress. Following egress, parasites migrate to and invade new host cells and the lytic cycle repeats, leading to tissue destruction (Black & Boothroyd, 2000). Actin plays an essential role in the parasite’s lytic cycle through function of the actin cytoskeleton and actomyosin motor complex. Despite this crucial role in apicomplexan biology, there has been difficulty visualising actin filaments in apicomplexan species (Bannister & Mitchell, 1995; Sahoo et al., 2006; Shaw & Tilney, 1999) and it has been suggested that as much as 98% of parasite actin is monomeric (G-actin) and not incorporated into filaments (F-actin) (Dobrowolski et al., 1997). This, along with structural differences found in actin of Apicomplexa (Pospich et al., 2017), led to the hypothesis that *Toxoplasma* F-actin has reduced stability, forming abnormally short filaments that are rapidly recycled to maintain essential cellular function (Pospich et al., 2017; Skillman et al., 2011). However, recent development of the actin-chromobody has allowed for the visualisation of long F-actin structures both in the parasite and extensive networks within the PV (Periz et al., 2017). Taken together, along with biochemical evidence, apicomplexan actin appears to be different to actin from other organisms (Frénal, Dubremetz, et al., 2017).

Actin turnover is regulated by actin binding proteins, of which *Toxoplasma* possesses a reduced repertoire, including ADF/cofilin, profilin, coronin and cyclase-associated protein (CAP) (Baum et al., 2006). Functional studies have shown ADF/cofilin and profilin to be essential for *Toxoplasma* progression through the lytic cycle, while coronin depletion had a modest impact on parasite invasion and egress (Mehta & Sibley, 2011; Plattner et al., 2008; Salamun et al., 2014). Apart from its localisation, the function of CAP in *Toxoplasma* has not been investigated. In the majority of eukaryotes, CAP is a highly conserved multidomain protein that regulates actin filament dynamics via two distinct mechanisms (Ono, 2013). CAP can bind and sequester G-actin using its CAP and X-linked retinitis pigmentosa 2 protein (CARP) domain and can regulate actin filament disassembly by promoting ADF/cofilin mediated severing using its helical folded domain (HFD). Through regulation of actin dynamics, by interacting with actin and other actin binding proteins, it has been shown that mouse CAP1 plays important roles in cell morphology, migration and endocytosis (Bertling et al., 2004). However, species belonging to the Apicomplexa phylum possess a truncated form of CAP, retaining only the conserved C-terminal G-actin-binding CARP domain (Hliscs et al., 2010). This conserved β-sheet domain has been shown to interact directly with monomeric actin, providing either sequestration or nucleotide exchange of G-actin in a concentration-dependent manner (Hliscs et al., 2010; Makkonen et al., 2013; Mattila et al., 2004). As such, apicomplexan CAP is hypothesised to regulate actin turnover solely through interaction with monomeric actin. Biochemical analysis of *Cryptosporidium parvum* CAP identified the formation of a dimer and G-actin sequestering activity, while *Plasmodium falciparum* CAP was shown to facilitate nucleotide exchange, loading ADP-actin monomers with ATP (Hliscs et al., 2010; Makkonen et al., 2013). A *Plasmodium berghei* CAP KO demonstrated that while PbCAP is dispensable for asexual blood stages *in vivo,* there is a complete defect in oocyst development in the insect vector which was overcome through complementation with *C. parvum* CAP (Hliscs et al., 2010). A PbCAP overexpression study revealed no defect in ookinete motility or oocyst development, however sporozoites displayed impaired gliding motility, invasion and salivary gland colonisation (Sato et al., 2016). Taken together, these results suggest that apicomplexan CAP may function as a dimer with the ability to interact with G-actin monomers to sequester them and/or facilitate their nucleotide exchange. While CAP has yet to be functionally characterised in *Toxoplasma,* Lorestani *et al* reported that CAP localises to the apex of intracellular parasites, a hub for events leading to egress and motility (Graindorge et al., 2016; Lorestani et al., 2012; Tosetti et al., 2019). Intriguingly, following host cell lysis, relocalisation of CAP to the parasite cytosol was observed (Lorestani et al., 2012). Furthermore, we have previously identified differential phosphorylation of TgCAP in parasites with a delayed egress phenotype (Treeck et al., 2014). The correlation between TgCAP redistribution and phosphorylation, following host cell lysis, hints at a potential role for TgCAP in actin regulation during rapid egress. Taken together, these results prompted us to characterise the role of TgCAP in *Toxoplasma* biology.

Here we show that TgCAP is produced in two distinctly localised isoforms through alternative translation initiation: a membrane bound isoform, localised to the apical tip (longCAP) and a cytosolically dispersed isoform (shortCAP). Conditional knockout of TgCAP, using a second generation DiCre strain, identified an important function of TgCAP in some, but not all, actin-dependent processes. Invasion, egress, motility, correct daughter cell orientation and dense granule trafficking were all perturbed in CAP depleted parasites, while apicoplast inheritance was not. This suggests different spatial requirements for CAP in actin turnover within the cell. Furthermore, the characteristic rosette organisation of parasites in the vacuole was completely lost, but synchronicity of division was unaffected. Strikingly, while we observe rapid protein transfer only between two adjacent cells in a vacuole, all parasites remain connected through a decentralised residual body, potentially explaining why synchrony of division is unaffected. In the mouse *in vivo* infection model, *CAP*-depleted type I RH parasites display normal lethality, while CAP depletion renders type II Pru parasites avirulent, with markedly reduced cyst formation. Furthermore, the cytoplasmic isoform of TgCAP was sufficient for the infection of mice and the formation of latent stages in the brain, indicating that the apically localised CAP isoform provides only a small fitness benefit under the conditions tested here.

## Results

### TgCAP contains two translational start sites which results in the production of two differentially localised protein isoforms

TgCAP was previously shown to localise to the apex of intracellular parasites and rapidly redistribute to the cytoplasm of extracellular parasites following host cell lysis (Lorestani et al., 2012). This suggested that TgCAP localisation may be influenced by post-translational modifications that enable CAP to regulate actin dynamics at different locations in the cell. TgCAP, and CAP from *Neospora* and *Hammondia* species contain a unique predicted N-terminal extension that is not present in other Apicomplexa, such as *Plasmodium* (Fig 1A). The extension contains two predicted palmitoylation sites and CAP was identified in an analysis of palmitoylated proteins in *Toxoplasma gondii* (Foe et al., 2015). Furthermore, two phosphorylation sites in the N-terminus of TgCAP are substantially phosphorylated upon ionophore-induced egress (Treeck et al., 2014). These phosphorylation events are dependent on the calcium-dependent kinase 3 (TgCDPK3) (Treeck et al., 2014), which has been shown to be important in mediating rapid exit from the host cell and is localised to the plasma membrane (Black et al., 2000; Garrison et al., 2012; Lourido et al., 2012; McCoy et al., 2012). Collectively, these observations allow for the possibility that re-localisation of CAP is important for egress and is mediated by dynamic post-translational modifications. To visualise TgCAP we expressed it as a GFP fusion. TgCAP-GFP localises to the apex of the parasite, as previously shown, but also to the cytoplasm of intracellular tachyzoites (Fig. 1B). We additionally demonstrated this dual localisation of TgCAP by C-terminally tagging the endogenous TgCAP locus with a HA epitope tag (Supplementary Fig. 1). To rule out any mis-localisation of the protein as a result of tagging, we expressed recombinant TgCAP, spanning residues 37 to 203, and generated antibodies against TgCAP which confirmed the dual localisation (Fig 1C). Western blot analysis of parasite lysates revealed the presence of two bands close to the expected size of TgCAP, which are expressed at a constant level, relative to the *Toxoplasma* loading control, across the first 24 hours following host cell invasion (Fig. 1D). The dual localisation of CAP could be due to the expression of two isoforms from a single gene, through use of alternate translational start sites, as previously observed for protein kinase G (Brown et al., 2017). Indeed, sequence comparison of TgCAP to PfCAP reveals a second in-frame methionine at position 37 in *Toxoplasma* that aligns with the PfCAP start methionine (Fig. 1A). Two additional methionine residues, M71 and M161, are present in the CAP primary sequence. However, these are unlikely used as translational start sites as their products would lead to a truncated and potentially inactive CARP domain.

**Figure 1:**
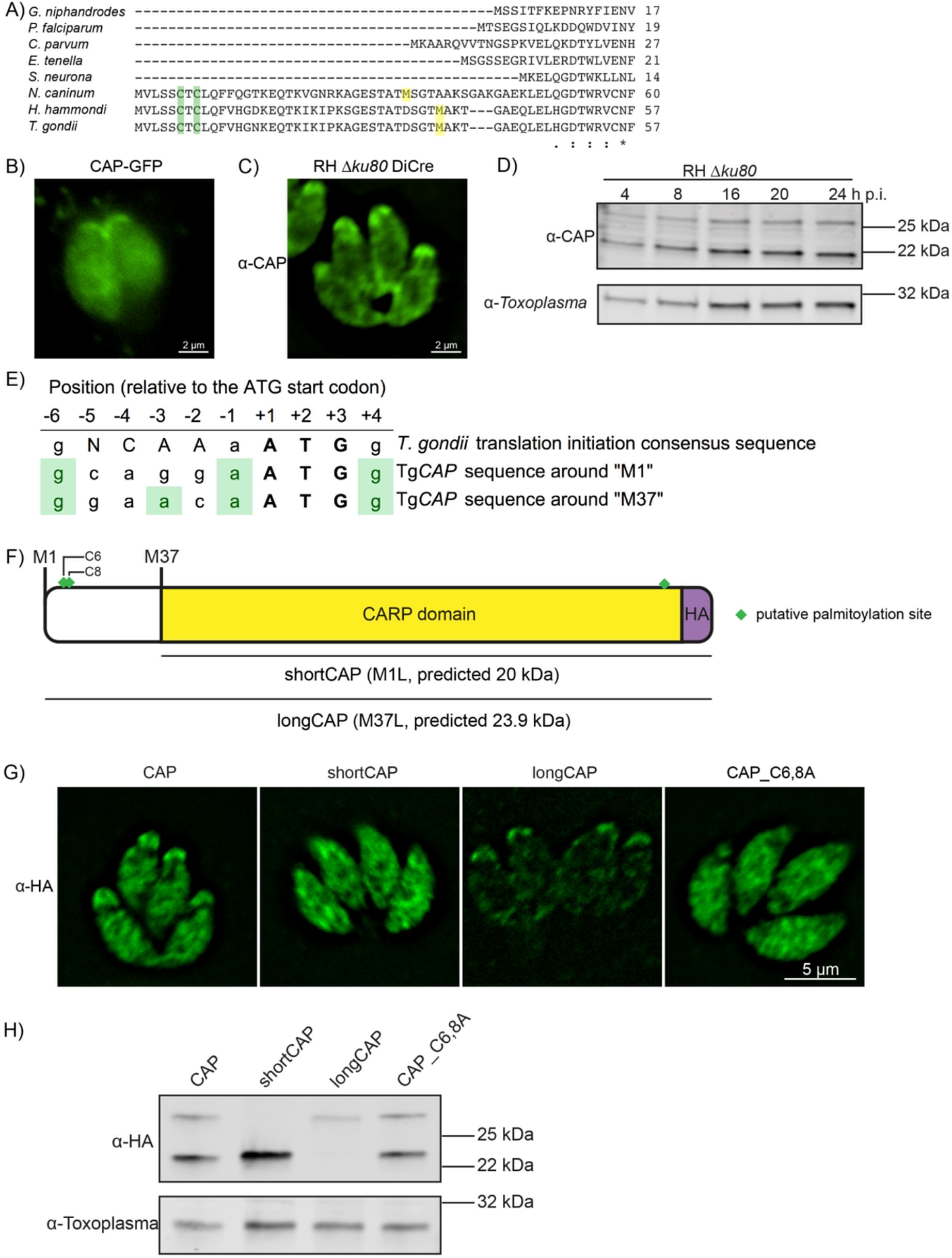
Alternative translational start sites lead to the generation of two different CAP isoforms. **(A)** Sequence alignment of the first 57 amino acid residues of TgCAP with that of other Apicomplexa. Green shading indicates cysteines which are putative palmitoylation sites, yellow shading indicates methionines which are putative alternative translational start sites. **(B)** Subcellular localisation of a CAP-GFP fusion. Scale bar, 2 μm. **(C)** Subcellular localisation of CAP by immunofluorescence assay (IFA) using rabbit anti-TgCAP antibodies. Scale bar, 2 μm. **(D)** Western blot of CAP expression levels over the first 24 hours following host cell invasion using anti-TgCAP antibodies. Anti-*Toxoplasma* antibodies were used as a loading control. **(E)** Alignment of the *Toxoplasma* consensus translation initiation (Kozak) sequence (Seeber, 1997) with the translation initiation sequences of *CAP’s* first (M1) and second (M37) putative translational start sites. Green shading indicates bases that correspond to the Kozak sequence. **(F)** Schematic of TgCAP with annotations for the two putative CAP isoforms and the mutation experiments performed to test their expression: Mutation of M1 to leucine (L) to produce shortCAP, and mutation of M37 to L producing longCAP. Green diamonds indicate putative palmitoylation sites. **(G)** IFA and **(H)** western blot of ectopic HA-tagged TgCAP isoforms and cysteine mutants (C6 and C8). Inclusion of a HA-tag makes the protein run more slowly than the untagged protein. Anti-*Toxoplasma* antibodies were used as a loading control in (H). Scale bar in (G), 5 μm.

Translational start sites in eukaryotic mRNAs are preceded by a translation initiation sequence (Kozak, 1987a, 1987b). Consensus translation initiation sequences have been determined for many different organisms and are known as the Kozak sequence (Nakagawa et al., 2008). In *Toxoplasma,* the Kozak sequence was elucidated by identifying nucleotides commonly found in highly abundant proteins, such as SAG1 (Seeber, 1997). This Kozak sequence contains an adenine at the −3 position, relative to the start ATG, and was identified as the most important factor in ribosomal recognition of the start ATG (Seeber, 1997). Absence of an adenine can result in ribosome “leaky scanning” and translation from an internal ATG. We therefore analysed the translation initiation sequence around CAP’s first (M1) and second (M37) putative translational start sites. The M1 translation initiation sequence conforms less with the *Toxoplasma* Kozak sequence than the sequence preceding M37; as the former is lacking an adenine at the −3 position (Fig. 1E). This suggested that alternative translation could lead to the generation of two CAP isoforms: longCAP, which is translated from the first start ATG, and shortCAP, which is translated from the second start ATG. To test this, we generated parasite strains that expressed either the WT sequence or variants where either the first (M1), or the second methionine (M37) of TgCAP was mutated to leucine, precluding their use as translational start sites (Fig. 1F). We used the endogenous promotor (i.e. 969 bp upstream of the first start ATG) and introduced the C-terminal HA-tagged TgCAP variants into the *UPRT* locus of the RH DiCreΔ*ku80*Δ*hxgprt* parasite strain. To determine whether longCAP and shortCAP show differential localisation, as predicted by the presence of putative palmitoylation sites in longCAP, we analysed their subcellular localisation using the HA-tag. While WT CAP parasites showed the expected dual localisation, mutants expressing shortCAP showed exclusively cytoplasmic staining while those expressing longCAP showed predominantly apical staining, with some further signal throughout the parasite (Fig. 1G). As longCAP contains two putative palmitoylation sites not present in the shortCAP sequence, we next evaluated whether palmitoylation was important for the apical localisation of the long CAP isoform. We mutated the two cysteines in the N-terminus to alanine residues (CAP_C6,8A). CAP_C6,8A appeared cytosolic with no detectable accumulation of CAP at the apical end of the parasites (Fig. 1G). Western blot analysis of the HA-tagged TgCAP variants confirmed that WT CAP is identified as two distinct protein bands, which correlate with the predicted size for longCAP and shortCAP (23.9 and 20 kDa). Parasites expressing shortCAP displayed only the lower molecular weight band, while longCAP-expressing parasites only showed the higher molecular weight band. We observed that the protein levels of longCAP appear reduced compared to its isoform in parasites expressing WT CAP, whereas the shortCAP isoform shows an increase (Fig. 1H). As expected, despite CAP_C6,8A not being detected at the apical end of the parasite, both isoforms were detected by Western blot at levels comparable to parasites expressing WT CAP. Collectively, these results show that in *Toxoplasma*, and potentially closely related coccidian parasites of *Hammondia* and *Neospora,* CAP is produced as two differentially localised isoforms using alternative translational start sites and the apical localisation of the long isoform is likely palmitoylation dependent.

### Generation of a more stable RH DiCre Δ*ku80* cell line

While attempting to generate a conditional knock out (cKO) of CAP using the DiCre strategy, we observed a frequent loss of one of the DiCre subunits, resulting in dysfunctional floxed CAP parasite strains that lacked the ability to excise CAP. This is possibly because both DiCre subunits are driven by identical 5’ and 3’ UTRs, allowing for potential recombination in the Δ*ku80* parental line, which possesses an increased efficiency of homologous recombination (Fox et al., 2009; Huynh & Carruthers, 2009). To prevent loss of DiCre subunits, we generated a new DiCre construct, DiCre_T2A, that expresses the two DiCre subunits from a single promotor using T2A skip peptides (Kim et al., 2011). To further minimise the potential for loss of DiCre, we placed a chloramphenicol acetyltransferase (CAT) selectable marker between the two subunits (Fig. 2A). This would lead to the production of the two separate Cre subunits and the CAT selectable marker. We inserted this construct, into the modified *KU80* locus of the RH Δ*ku80*Δ*hxgprt* strain (Huynh & Carruthers, 2009) using CRISPR/Cas9. To test whether expression of the DiCre subunits in the resulting line, RH DiCre_T2A Δ*ku80*Δ*hxgprt*, is stable over time, we integrated the loxP-KillerRed-loxP-YFP reporter construct used in Andenmatten *et al.,* into the *UPRT* locus (Andenmatten et al., 2013). As expected, non-treated parasites express KillerRed which, upon RAP treatment, is excised and leads to expression of YFP (Fig. 2B). As extracellular stress can lead to increased loss of DiCre activity in the original DiCre line (M. Meissner, personal communication, 02.2019), we subjected the new DiCre parasite line (RH DiCre_T2A Δ*ku80*Δ*hxgprt*) to frequent extracellular stress over the course of 65 days. On average, parasites were passaged every 2.3 days, leaving parasites extracellular for ~32 hours in the presence or absence of continuous chloramphenicol selection. We also simultaneously passaged the original DiCre line (Andenmatten et al., 2013) under standard, non-stressing, culturing conditions. The new DiCre_T2A line excision efficiency varied between 98 and 99% for replicates on day 1 and was maintained throughout the experiment with a maximal loss of 3% of excision efficiency, irrespective of the presence or absence of chloramphenicol selection, by day 65 (Fig. 2C). In contrast, the original DiCre line, cultured under standard non-stress conditions, lost 42% of excision capacity by day 65 (Supplementary Fig. 2), although this was only done as a single replicate. This shows that the second generation DiCre line, RH DiCre_T2A Δ*ku80*Δ*hxgprt*, retains high excision capacity over long periods of time, even when exposed to extracellular stress.

**Figure 2:**
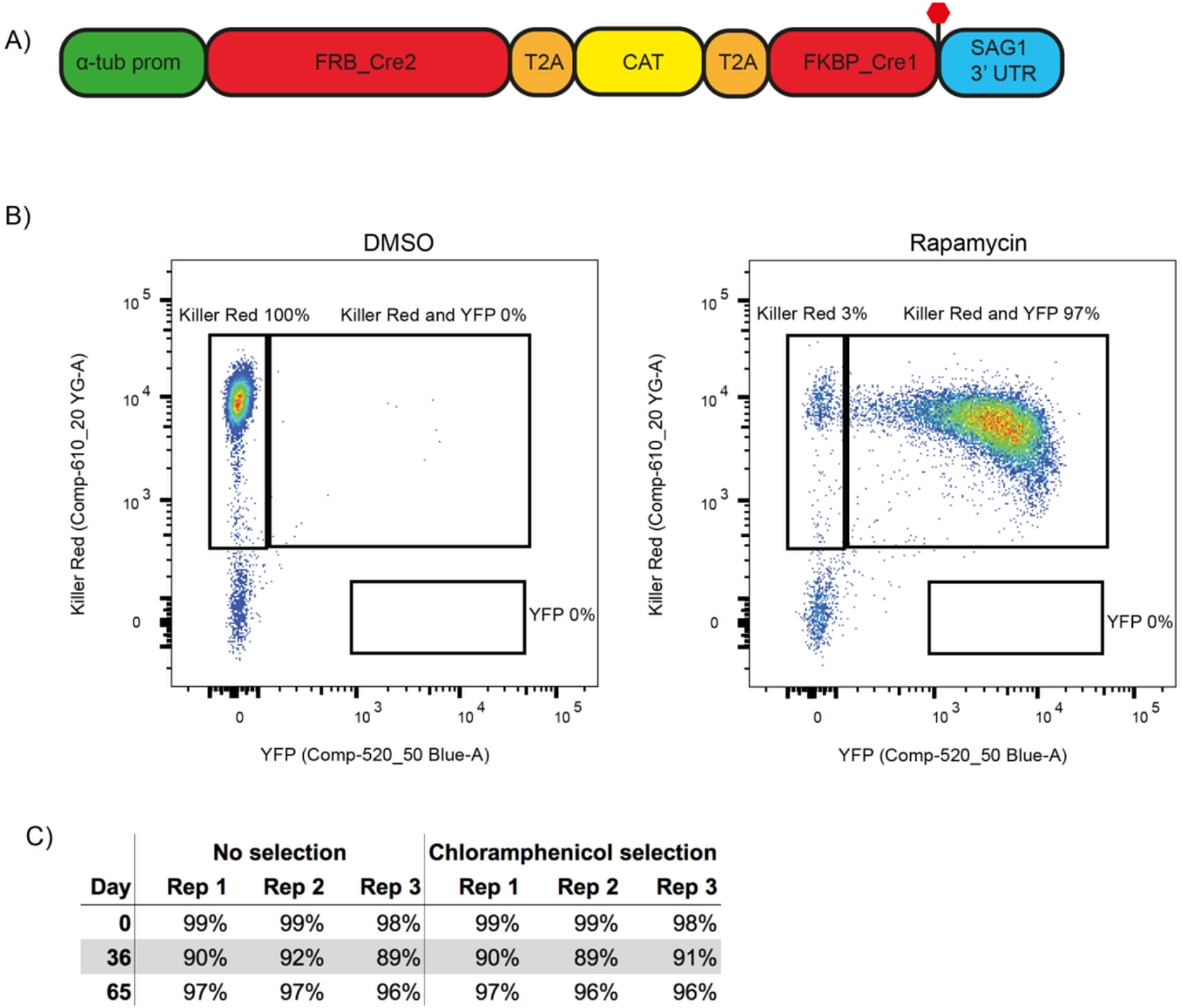
A second generation RH Δ*ku80* DiCre_T2A parasite strain stably expresses DiCre. **(A)** Schematic of the DiCre_T2A expression construct. The chloramphenicol resistance cassette (CAT) is flanked by T2A skip peptides. The two Cre subunits (FRB_Cre2 and FKBP_Cre1) are located on either side of the T2A::CAT::T2A cassette. The fusion protein is driven by the alpha-tubulin promotor with a *SAG1* 3’ UTR. The red hexagon indicates the position of the stop-codon. **(B)** Flow cytometry analysis to determine excision efficiency of the RH DiCre_T2A Δ*ku80* line following 65 days of frequent extracellular stress. Excision is determined by a shift from Killer Red^(+)^ to Killer Red^(+)^ and YFP^(+)^ expression. Parasites were analysed 22 h after induction with 50 nM rapamycin (RAP) for 4 h. Due to analysing 22 h after induction of excision, parasites still have residual KillerRed signal. **(C)** Table summarizing the excision efficiency of RAP treated RH DiCre_T2A Δ*ku80* parasites over time in the presence or absence of chloramphenicol selection. “Day” refers to the number of days in cell culture while “rep” corresponds to biological replicates.

### TgCAP is important but not essential for *in vitro* growth and deletion can be largely restored by the short cytoplasmic isoform, but only partially by the membrane bound isoform

To investigate CAP function, we generated a conditional knock out (cKO) of *CAP* using the DiCre strategy. Here, *CAP* with a C-terminal HA tag is flanked by two loxP sites, that recombine upon dimerisation of two split-Cre subunits, a process mediated by the small molecule rapamycin (RAP) (Andenmatten et al., 2013). To create the *CAP* conditional knockout line, we integrated a floxed, recodonised and HA-tagged CAP cDNA sequence into the endogenous locus of the RH DiCreΔ*ku80*Δ*hxgprt* line (Fig. 3A). Correct integration into the locus was confirmed by PCR (Fig. 3B). However, due to the DiCre issues detailed above, we were not able to successfully induce DiCre-mediated excision of the *CAP* gene. As the DiCre_T2A strategy demonstrated consistently high excision rates over time, during our testing with a reporter construct, we integrated the DiCre_T2A construct into the *KU80* locus of the non-excising floxed *CAP* parasite strain. This generated the parasite line RH DiCre_T2A DiCreΔ*ku80*Δ*hxgprt*_*LoxPCAP-HA,* called LoxPCAP hereafter. As expected, LoxPCAP displayed dual localisation by IFA (Fig. 3C). RAP treatment resulted in a complete loss of CAP (ΔCAP), as shown by IFA (Fig. 3C) and Western blot (Fig. 3D).

**Figure 3:**
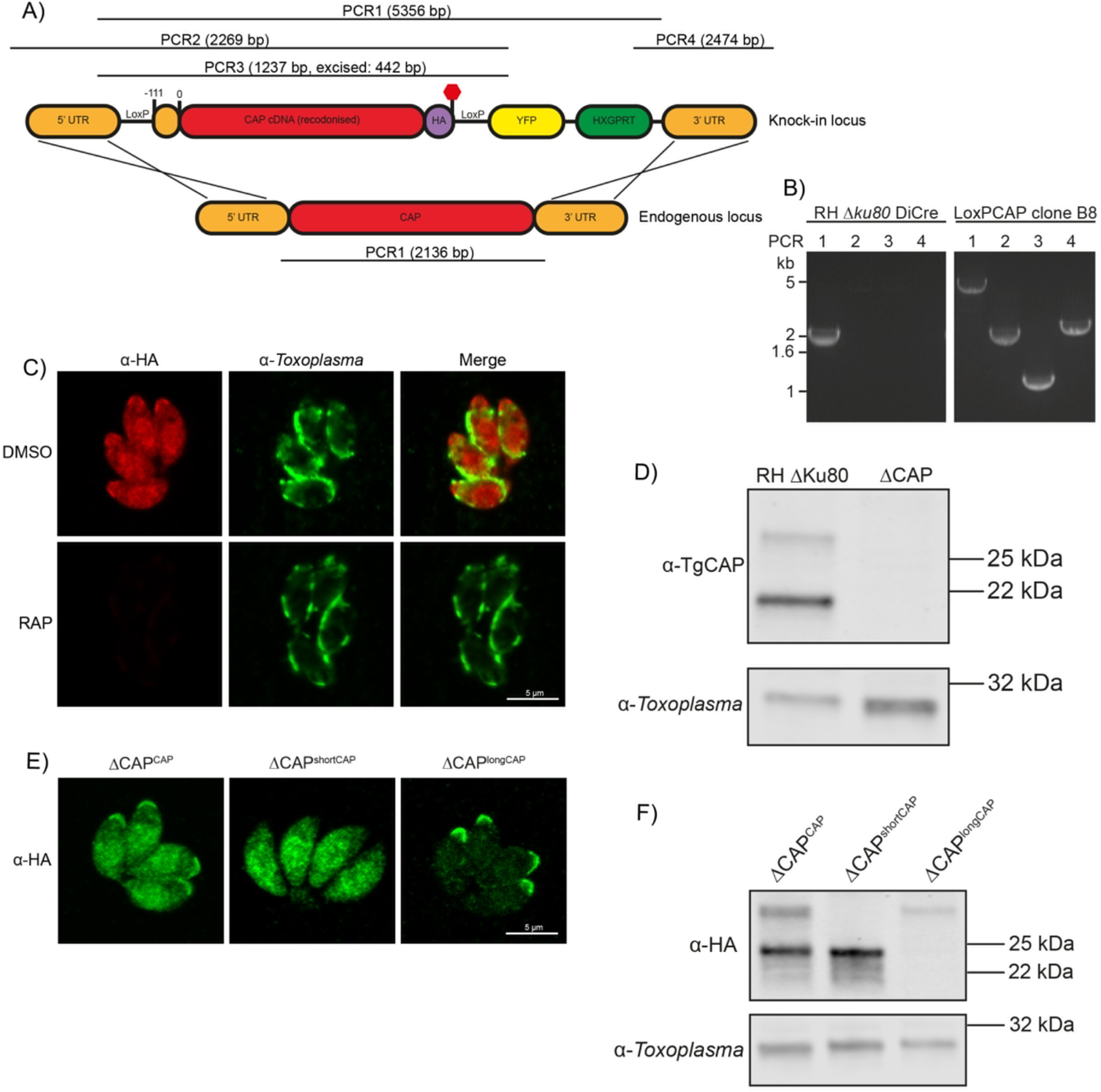
Generation of CAP conditional knockout and complementation strains. **(A)** Schematic of the CAP conditional knock out strategy using double homologous integration. The position of the 5’ LoxP site in the CAP promotor is indicated as well as the predicted sizes of the PCR amplicons. **(B)** Agarose gel showing the expected PCR products for correct integration at the endogenous locus. **(C)** IFA 46 hr after treatment with DMSO or 50 nM rapamycin for 4 h. Scale bar, 5 μm. **(D)** Western blot showing absence of CAP protein in the cloned ΔCAP parasites using anti-TgCAP antibodies. Anti-*Toxoplasma* antibodies were used as a loading control. **(E)** Subcellular localisation and **(F)** Western blot of ectopic HA-tagged TgCAP isoforms, in the ΔCAP background. Inclusion of a HA-tag makes the protein run more slowly than the untagged protein. IFA images have been individually contrast adjusted to aid in visualising protein localisation. Scale bar in (E), 5 μm. Anti-*Toxoplasma* antibodies were used as a loading control in (F).

To assess the requirements of CAP and its isoforms for various *Toxoplasma* functions, we complemented LoxPCAP parasites with either wildtype, the short or the long CAP isoform by integration of HA-tagged variants into the *UPRT* locus to generate merodiploid lines (named LoxPCAP^CAP^, LoxPCAP^shortCAP^ and LoxPCAP^longCAP^, respectively). We then excised the endogenous CAP copy by RAP treatment. Clones were subsequently obtained by limiting dilution and excision verified by PCR (Supplementary Fig. 3).

This resulted in parasite strains that express only the WT complemented form (ΔCAP^CAP^), the short complemented form (ΔCAP^shortCAP^) or the long complemented form (ΔCAP^longCAP^). We confirmed the differential localisation and translation of these isoforms by IFA (Fig. 3E) and Western blot (Fig. 3F). No differences in the protein levels of the two CAP isoforms could be observed in the CAP complemented strain (ΔCAP^CAP^) over the first 24 hours following host cell invasion, relative to the loading control (Supplementary Fig. 4).

To reliably quantify the contribution of CAP, and its two isoforms independently, to the lytic cycle, we performed a competition assay in which growth of ΔCAP^CAP^ was compared to ΔCAP, ΔCAP^shortCAP^ and ΔCAP^longCAP^. To do so, we integrated an mCherry expressing cassette into the Δ*ku80* locus, replacing the DiCre_T2A cassette in each of these lines. Note, the DiCre_T2A cassette was no longer required because CAP had already been excised in these parasites. After 15 days in growth competition with ΔCAP^CAP^, ΔCAp parasites were largely depleted (>97.3%) from the population, while ΔCAP^shortCAP^ showed a reduction of only 4%. In contrast, ΔCAP^longCAP^ showed an intermediate level of depletion (33.3%). These phenotypes were exaggerated after 30 days in culture; ΔCAP parasites were largely depleted (>99.9%), ΔCAP^shortCAP^ showed a reduction of 12.4%, and ΔCAP^longCAP^ growth was reduced by 43.1% relative to ΔCAP^CAP^ (Fig. 4A). Collectively, these findings demonstrate that CAP plays an important but non-essential role in cell culture and that its function can be largely restored by the short CAP isoform but only partially by the long isoform.

**Figure 4:**
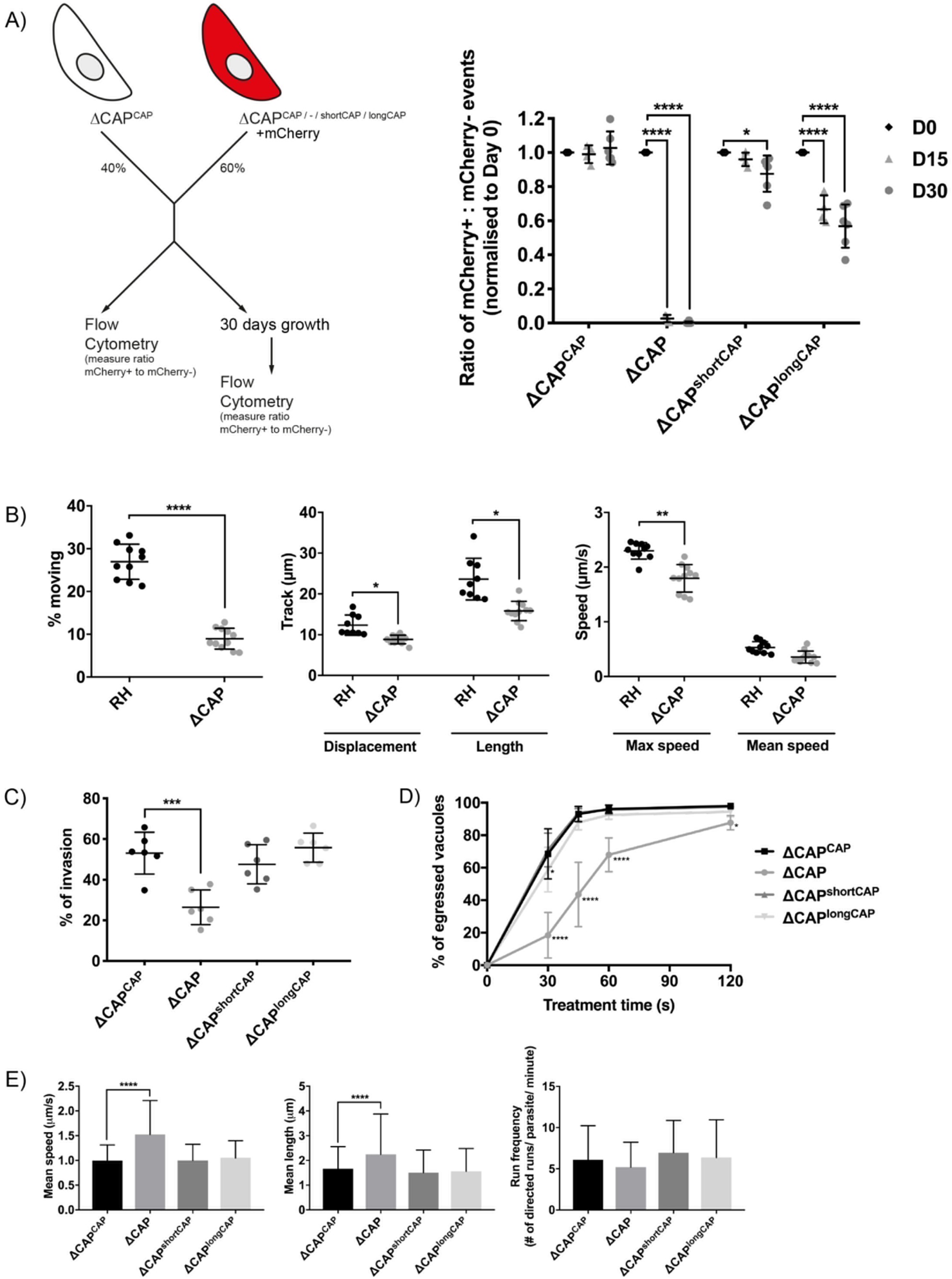
CAP plays an important but not essential role during the lytic cycle in cell culture. **(A)** Overview of the flow cytometry competition assay (left). Competition assays of mCherry-expressing ΔCAP and CAP complementation lines with non-fluorescent WT-complemented parasites (right). The ratio of mCherry^(+)^ to mCherry^(-)^ parasites was analysed by flow cytometry at day 0, 15 and 30. Data are represented as mean ± s.d. (D0 and D30, *n*=6. D15, *n*=4). Two-way ANOVA followed by a multiple comparison Sidak’s test was used to compare means between time points. **(B)** 3D Matrigel-based motility assays performed in the absence of inducers of motility. Results are expressed as mean ± s.d. (*n*=4). Each data point corresponds to a single technical replicate from one of four independent biological replicates, on which significance was assessed using an unpaired *t*-test. **(C)** Invasion assay comparing ΔCAP to the complemented lines. Data are represented as mean ± s.d. (*n*=3). One-way ANOVA followed by Dunnett’s test was used to compare means to the ΔCAP^CAP^ mean **(D)** Egress assay. Graph shows number of egressed vacuoles in response to BIPPO over time. Data are represented as mean ± s.d. (*n*=3). Two-way ANOVA followed by Dunnett’s test was used to compare means to the ΔCAP^CAP^ mean. Stated significance is in comparison to ΔCAP^CAP^. **(E)** Dense granule trafficking assay. ΔCAP and CAP complemented lines were transiently transfected with SAGIΔGPI-mCherry that allows visualisation of dense granules. The length and speed of directed runs were recorded using fluorescence microscopy and analysed using ImageJ. Data are represented as mean ± s.d. (*n*=3). One-way ANOVA followed by Dunnett’s test was used to compare means to the ΔCAP^CAP^ mean.

### TgCAP contributes to motility, invasion, egress and dense granule trafficking

While the competition assays highlight the importance of CAP during the *in vitro* lytic cycle, they do not clarify which step of the cycle is affected. To test whether the growth differences between the lines were merely a result of differences in parasite replication rates, we counted the parasites per vacuole for each parasite line. No significant differences were observed (Supplementary Fig. 5). As CAP is a predicted actin regulator, we next focused our phenotypic analysis on *Toxoplasma* processes for which actin is known to be important, such as motility, egress, invasion and dense granule trafficking (Heaslip et al., 2016; Periz et al., 2017; Whitelaw et al., 2017).

In the absence of CAP (ΔCAP) significantly fewer parasites are able to initiate gliding motility in 3D motility assays (Leung et al., 2014) compared to the RH strain (Fig. 4B). In the motile population, trajectory displacement, trajectory length and maximum achieved speeds were all significantly reduced, although mean speed was not significantly different (Fig. 4B). To determine which isoform(s) contribute to parasite 3D motility, the motility parameters of ΔCAP^CAP^, ΔCAP^shortCAP^ and ΔCAP^longCAP^ were compared. Both ΔCAP^shortCAP^ and ΔCAP^longCAP^ parasites showed similar levels of motility initiation, track displacement, track lengths and speed compared to ΔCAP^CAP^ (Supplementary Fig. 6). These data indicate that CAP plays a role in initiation of motility and in controlling speed and track length once motile. Complementation with single isoforms shows that initiation of motility can be rescued by either CAP isoform and, once motile, either can maintain speed and track length.

As invasion and egress of host cells rely on active motility, we next compared invasion efficiency of ΔCAP and the different complementation lines. ΔCAP showed a significant reduction of invasion capacity (50.2% reduction compared to WT complemented lines), which was restored by both the short and long isoforms (Fig. 4C). We also performed egress assays in the presence of BIPPO, which results in a strong calcium response in *Toxoplasma* parasites causing synchronised and rapid egress from host cells (Howard et al., 2015). ΔCAP parasites showed a substantial delay in egress from host cells at 30 s after induction (73.1% less egress in ΔCAP compared to ΔCAP^CAP^), while at 2 minutes the majority of parasites have egressed (10.6% less egress in ΔCAP compared to ΔCAP^CAP^) (Fig. 4D). This defect was fully restored in ΔCAP^shortCAP^ parasites, while ΔCAP^longCAP^ showed slightly lower levels of egress after 30 s of treatment but by 60 s were indistinguishable from WT or short CAP complemented lines. We also observed live egress events in which ΔCAP, following host cell egress, showed decreased movement away from the host cell when compared to ΔCAP^CAP^, ΔCAP^shortCAP^ and ΔCAP^longCAP^ (Videos 1-4).

A recent study revealed that directed dense granule transport is dependent on filamentous actin (Heaslip et al., 2016). Therefore, dense granule trafficking was assessed in ΔCAP and the three complemented lines. We measured the run frequency (# of directed runs/parasite/minute), run length and velocity of directed motions. (Fig. 4E). Both the mean run length and the mean velocity were significantly increased in ΔCAP (34.8% and 52.9%, respectively), but not in either of the complements. However, the frequency of directed runs was reduced in ΔCAP (14.8%), but did not reach significance. These results indicate that both CAP isoforms play a supporting role in dense granule trafficking and that, upon CAP deletion, dense granules are trafficked further distances at higher speed. Collectively these data show that CAP plays a role in the actin-dependent processes described above and that either isoform is able to fulfil these CAP functions.

### ΔCAP parasites display a defect in efficient cell-cell communication

Actin, formin 3 and myosin I and J have been shown to be important for both parasite rosette organisation and cell-cell communication, as assessed by measuring the rapid transfer of reporter proteins between individual parasites in a vacuole (Frénal, Jacot, et al., 2017; Periz et al., 2017; Tosetti et al., 2019). Upon CAP deletion, we observed a complete loss of the characteristic rosette formation normally seen in *Toxoplasma*. This aberrant phenotype was fully rescued upon complementation with either WT CAP or the short CAP isoform (Fig. 5A). ΔCAP^longCAP^ displayed partial restoration with 44.9% of parasites forming phenotypically normal rosettes while the remainder were disorganised within the vacuole. This was also shown by scanning electron images of infected human fibroblasts in which the host cell and the vacuole membrane was removed (“unroofed”) as previously described (Magno et al., 2005) (Supplementary Fig. 7).

**Figure 5:**
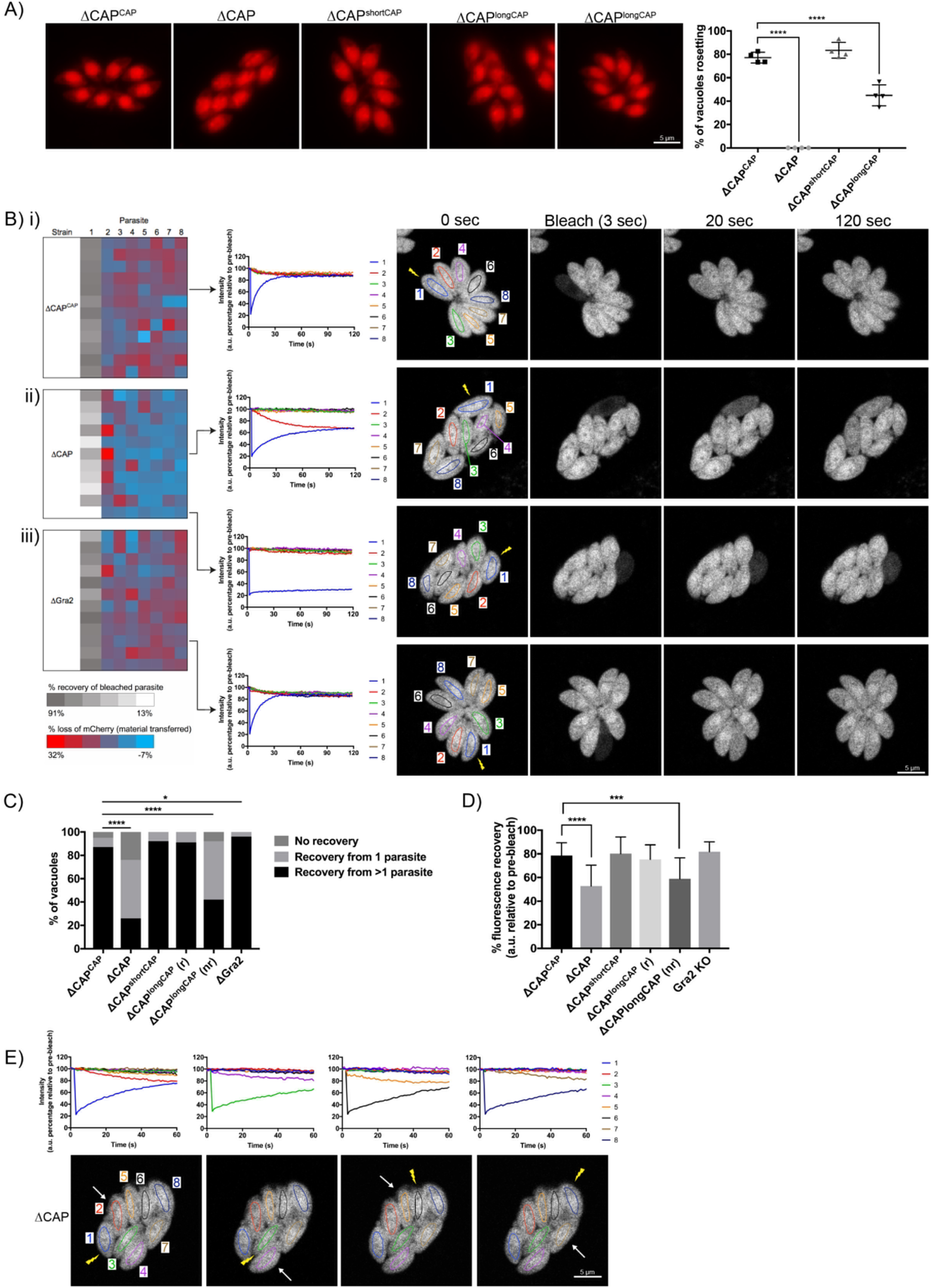
CAP is important for rosetting and rapid cell-cell communication. **(A)** Representative fluorescence images of mCherry expressing ΔCAP parasites and complemented isoforms (left). Quantification of rosetting vs. non-rosetting parasites (right). Data are represented as mean ± s.d. (*n*=2). One-way ANOVA followed by Dunnett’s test was used to compare means to the ΔCAP^CAP^ mean. **(B-E)** FRAP experiments to measure transfer of mCherry between individual parasites in a vacuole. **(B)** Heatmaps showing the percentage change in fluorescence of individual parasites in **(i)** CAP complemented, **(ii)** ΔCAP and **(iii)** Δ*gra2* parasite lines. A recovery plot and image for representative vacuoles are included. Regions of interest are numbered; the bleached parasite is “1”. Numbers are allocated based on proximity to the bleached parasite. The yellow lightning bolt indicates which parasite was photobleached. Data are representative of two independent experiments. **(C, D)** Graphs quantifying the type (C) and amount (D) of recovery for all parasite lines. Data are represented as mean ± s.d. (ΔCAP and ΔCAP^CAP^ *n*=4, all other lines, *n*=2). Statistical significance was assessed by either Chi-square test (D) or one-way ANOVA followed by Dunnett’s test to compare means to the ΔCAP^CAP^ mean (C). **(E)** FRAP analysis of ΔCAP parasites. The images and graphs represent sequential photobleaching and recovery measurements of individual parasites within the vacuole. The yellow lightning bolt indicates which parasite was photobleached. The white arrow identifies the parasite from which the majority of recovery was observed. All scale bars, 5 μm.

It was previously shown that mutant parasite lines which lost rosetting capacity also lost the **r**apid **t**ransfer **o**f **r**eporter **p**roteins (called RTORP hereafter) between them. This has been established by photobleaching of individual parasites in a vacuole, which express a fluorescent reporter protein. Under normal conditions, all parasites in a vacuole contribute to the fluorescence recovery of the photobleached parasite (Frénal, Jacot, et al., 2017; Periz et al., 2017; Foe et al., 2018; Tosetti et al., 2019). Such transfer of reporter proteins has previously been used as a readout for cell-cell communication and, by extension, parasite connectivity (Frénal, Jacot, et al., 2017; Periz et al., 2017). To examine whether loss of cell-cell communication also accompanied the disrupted rosetting observed in ΔCAP parasites, we performed fluorescence recovery after photobleaching (FRAP) experiments on ΔCAP and its complementation lines. We chose vacuoles that contained 8 parasites/ vacuole as at this stage parasites organise in rosettes and individual cells can be easily monitored. We bleached one parasite in the vacuole and recorded both the recovery of fluorescence in the bleached parasite and the fluorescence levels of all other parasites in the vacuole. As expected, the WT-complemented ΔCAP^CAP^ photobleached parasites display rapid recovery of fluorescence to which, in most cases, all parasites in the vacuole appear to contribute (Fig. 5Bi, 5C and Video 5). Conversely, ΔCAP photobleached parasites were supported in their rapid fluorescence recovery predominantly by just one other parasite, usually the parasite closest to the bleached cell, resulting in slow recovery (Fig. 5Bii, 5C and Video 6). In some cases, no recovery was observed in ΔCAP parasites.

Loss of rapid protein transfer between parasites could be explained by a structural disruption of inter-parasite connections provided by the residual body. However, in addition to the residual body, an intravacuolar network (IVN) of tubule-like structures is present in the parasitophorous vacuole. Because of its tubular structure, it could also be involved in cell-cell communication. To determine whether the observed defect in the RTORP was dependent on the presence of the IVN, we performed a FRAP assay in RH Δ*ku80*Δ*gra2* parasites (Rommereim et al., 2016) where the IVN fails to form but parasites still organise in rosettes. The results show that RTORP between parasites was not negatively affected (Fig. 5Biii and Video 7). The IVN, therefore, is unlikely involved in cell-cell communication.

To test if CAP deletion leads to a defect in cell-cell communication directly, or whether this is a direct consequence of the inability to form rosettes, we tested whether the RTORP varied between rosetting (r) and non-rosetting parasites (nr) of the ΔCAP^longCAP^ line. These experiments revealed that if ΔCAP^longCAP^ parasites rosette, they show normal RTORP between all cells in the vacuole, while non-rosetting parasites display the same defect as the ΔCAP line, with fluorescence recovery from only one other parasite in the vacuole (Fig. 5C). Interestingly, Δ*gra2* vacuoles showed a slight increase in the frequency of photobleach recovery from multiple parasites, but we have not further investigated this phenomenon here. The connectivity of parasites was also assessed based on the percentage of recovery after photobleaching. As expected, ΔCAP and non-rosetting ΔCAP^longCAP^ photobleached parasites recovered significantly less fluorescence than their rosetting counterparts (Fig. 5D).

Next, we sequentially photobleached individual ΔCAP parasites in a vacuole and identified which parasites appear to be physically connected based on the transfer of mCherry (Fig. 5E). This revealed that only parasite pairs in close proximity are rapidly communicating. Collectively the results show that CAP deletion leads to a loss of both rosetting and the rapid transfer of reporter proteins between more than two parasites. Furthermore, RTORP appears to be dependent on parasite rosette organisation, but not on presence of the IVN.

### ΔCAP parasites have a defect in daughter cell orientation but not in synchronised division or apicoplast inheritance

Previously, cell-cell communication has been hypothesised to control synchronous division between parasites within the same vacuole (Frénal, Jacot, et al., 2017). To determine whether ΔCAP parasites display phenotypes previously observed for mutants with defective cell-cell communication, we used IMC3 antibodies to visualise synchronicity of forming daughter cells (Fig. 6A). Surprisingly, ΔCAP vacuoles exhibited synchronous division with no significant difference to the WT complemented line (Fig. 6B). IMC3 staining was also used to assess daughter cell orientation. While in WT CAP complemented parasites 94.6% of daughter cells grew in the same orientation, a significant defect was observed in ΔCAP and ΔCAP^longCAP^ strains; only 42.9% (ΔCAP) and 67.7% (ΔCAP^longCAP^) of daughter cells orientated in the same direction. ΔCAP^shortCAP^ parasites were able to largely overcome this defect, with 89.5% of daughter cells growing in the same orientation (Fig. 6C). Such improper orientation of daughter cells following budding raises the possibility of improper organelle segregation too. To test this, we looked at apicoplast segregation, another actin-dependent process (Andenmatten et al., 2013; Jacot et al., 2013). Using streptavidin as a marker for the apicoplast, we identified no significant differences in apicoplast inheritance rates between ΔCAP^CAP^ and ΔCAP strains (Supplementary Fig. 8). These data show that despite a loss of rosette organisation and RTORP between parasites in a ΔCAP vacuole, as well as disordered daughter cell orientation, parasite division remains synchronous. Furthermore, while CAP is supporting many actin-dependent processes in *Toxoplasma,* it appears dispensable for apicoplast division.

**Figure 6:**
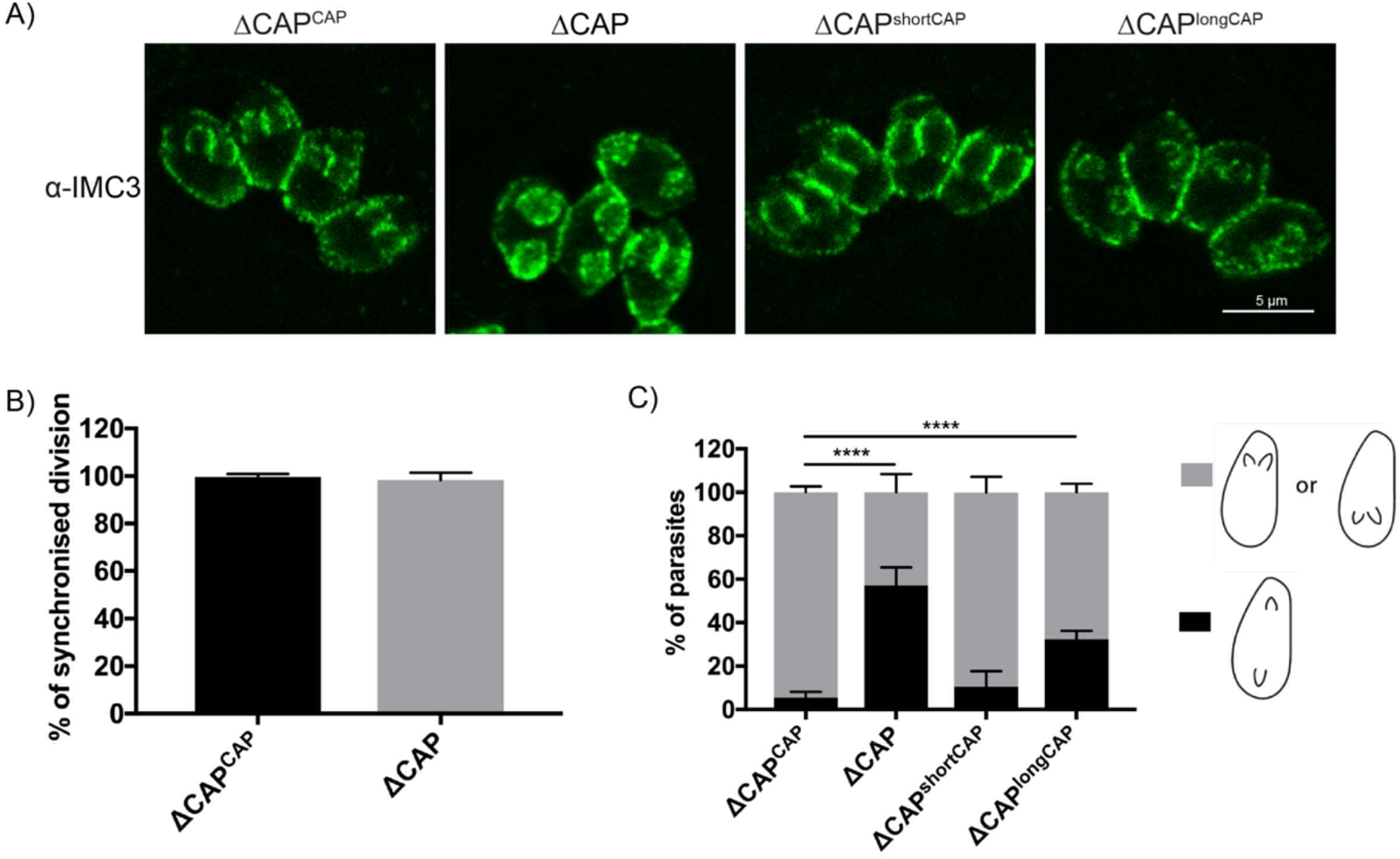
CAP is important for daughter cell orientation, but not synchronous division. **(A)** Parasites stained with anti-IMC 3 antibodies to visualise daughter cell orientation and division. Scale bar, 5 μm. **(B)** Quantification of synchronicity of division in parasite vacuoles using daughter cell staining from (A) reveals no defect in the synchronicity of ΔCAP parasites. Data are represented as mean ± s.d. (*n*=3). Significance was assessed using an unpaired two-tailed *t*-test. **(C)** Quantification of daughter cell orientation in parasite vacuoles reveals a significant defect in the daughter cell orientation of ΔCAP parasites. Data are represented as mean ± s.d. (*n*=3). One-way ANOVA followed by Dunnett’s test was used to compare means to the ΔCAP^CAP^ mean.

### CAP deletion leads to the formation of a decentralised residual body in which all parasites remain connected, despite loss of RTORP

The observed synchronicity of division in the ΔCAP strain, despite loss of RTORP, was unexpected. It suggested that synchronicity is either independent of the residual body, or that ΔCAP parasites actually maintain connections that allow flow of proteins or metabolites to synchronise divisions. To investigate this, we established a connectivity map of parasites in a vacuole using correlative light and electron microscopy (CLEM). This allowed us to first analyse connectivity of parasites based on FRAP analysis, and then reconstruct a 3D electron microscopy image of the parasites and their connections by focused ion beam scanning electron microscopy (FIB SEM).

In the ΔCAP^CAP^ strain, as expected for a WT complemented line, all parasites in the vacuole contribute to recovery of a photobleached parasite (Fig. 7A) and a normal residual body is formed, connecting all parasites in the vacuole (Fig. 7B and Video 8), with one tubular extension extending away from the residual body with no apparent connections at the distal end. In contrast, for the ΔCAP strain, fluorescence recovery of the photobleached parasite was predominantly observed from those cells in close proximity (Fig. 7C). To investigate how these parasites are connected, we analysed FIB SEM images obtained from the photobleached vacuole in Fig. 7C. This revealed membrane bound tubular connections of approximately 300 nm thickness between parasites, despite the aberrant transfer of mCherry and loss of rosetting (Fig. 7Di and ii). Following the lumen of the connections across 3 dimensions demonstrates that all parasites in the vacuole are connected by these tubular connections, likely representing a decentralised residual body that forms as a result of an inability to keep the posterior ends of the parasites in close proximity (Fig.7D.iii, Supplementary Fig. 9 and Video 9). Correlation of the FRAP data with the FIB SEM images showed that rapid transfer of mCherry was always between parasites in close proximity at their basal ends.

**Figure 7:**
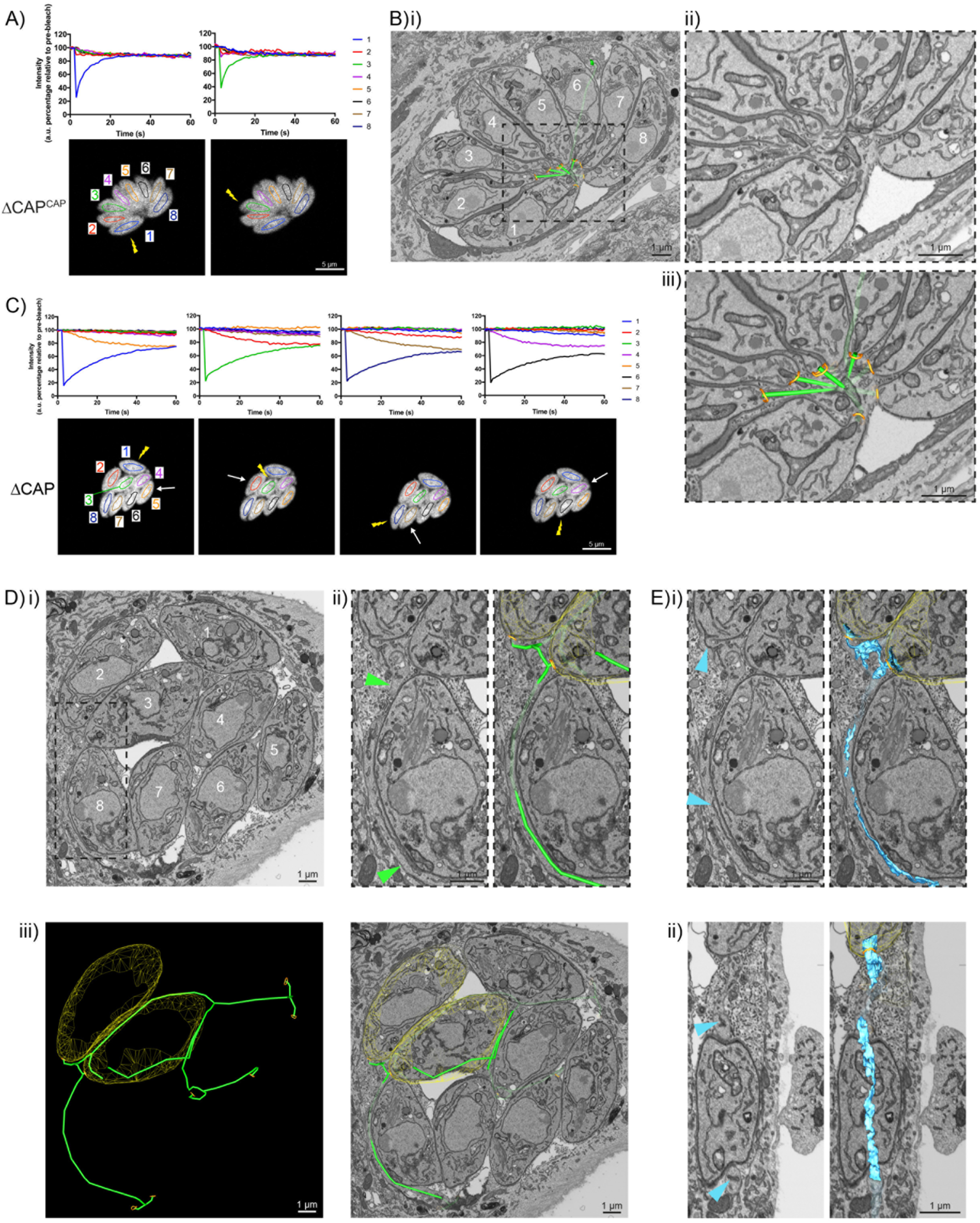
CAP KO parasites are still connected by a decentralised residual body. **(A)** FRAP analysis of ΔCAP^CAP^ parasites. The images and graphs represent sequential photobleaching and recovery measurements of individual parasites within the vacuole. **(B) (i)** FIB SEM of the vacuole from (A) with a 3D model highlighting the residual body (green skeleton representing how the approximate centre can be traced through to the basal poles) and parasite openings at the basal pole (orange ring). Note that the illustrated connections display the distance between the parasite’s posterior ends, and do not represent tubular connections between them. Numbering of parasites is consistent with (A). **(ii)** Zoomed image of the residual body from (i), **(iii)** zoomed image of the residual body from (i) overlaid with the 3D model. **(C)** FRAP analysis of ΔCAP parasites. The images and graphs represent sequential photobleaching and recovery measurements of individual parasites within the vacuole. **(D) (i)** FIB SEM of the vacuole from (A), numbering of parasites is consistent with (C). **(ii)** Zoomed image of the boxed region from (i). Green arrows indicate the residual body (left panel). A 3D model highlights the residual body (green skeleton drawn through the lumen of the connections between parasites), parasite openings at the basal pole (orange ring) (right panel). **(iii)** A 3D model of selected features in the vacuole from (C), highlighting the residual body (green), parasite openings at the basal pole (orange ring) and coarse segmentation of two of the parasites (yellow) (left panel). This model is shown with an orthoslice from the FIB SEM volume (right panel). **(E)** Zoomed images of the boxed region from (Di). Blue arrows indicate a putative ER structure (left). A 3D model highlights this putative ER (blue) and parasite openings at the basal pole (orange ring) (right). **(i)** View facing a Z orthoslice. **(ii)** View facing an X orthoslice. The yellow lightning bolt indicates which parasite was photobleached and the white arrow identifies the parasite from which recovery was observed (A,C). Scale bar, 5 μm (A, C) or 1 μm (B, D, E).

While ΔCAP parasites appear to be connected via a decentralised residual body, it could be that transfer of material through these connections is limited by physical barriers, such as the mitochondria which have previously been observed in the residual body (Frénal, Jacot, et al., 2017). Close examination of the residual body connections showed only one such connection contained two mitochondria and a constriction of the decentralised residual body lumen to ~50 nm, while all other connections appeared free of large physical barriers, indicating that this is unlikely an explanation for the lack of RTORP between parasites. However, we observed a complex network of tubules and sheet-like structures, likely representing the endoplasmic reticulum, in the connections (Fig. 7E) (Puhka et al., 2012; Schroeder et al., 2019; Tomavo et al., 2013; West et al., 2011). One of these tubular structures, tracked in three dimensions, was shown to enter multiple parasites via the basal pole, suggesting a possible continuum between them, which was also observed in then WT CAP complemented parasites. This has not been further examined here but could contribute to exchange of material between parasites.

Collectively, while CAP is important for rosette organisation of parasites, it is not essential for forming and sustaining a residual body with connectivity to all parasites in the vacuole. Furthermore, it shows that the RTORP is not an indicator of parasite connectivity.

### Deletion of CAP results in completely avirulent type II parasites, but not in the type I RH strain

The short CAP isoform complements most phenotypes in cell culture while the long CAP isoform, in most cases, only shows a partial rescue. This raises the question about the evolutionary roles of the two different isoforms. To better discriminate the functions of the short and longCAP isoforms, we wanted to examine their respective roles in natural infections, where parasites encounter a number of additional stresses, including shear stress and the immune system. Accordingly, we addressed the essentiality of CAP and its isoforms in mouse infections. We hypothesised that if both isoforms are essential for parasite survival in a natural host, loss of either of the two isoforms would manifest in a fitness cost. First, we injected male C57BL/6 mice with ~25 ΔCAP or ΔCAP^CAP^ tachyzoites of the virulent RH strain, and monitored them over the course of 10 days. In both instances mice began to succumb to infection after 9 days, indicating that, despite the motility and rosetting phenotypes, CAP depletion in a type I RH background still results in high virulence in mice (Fig. 8A). As RH parasites have frequently been associated with hypervirulence and do not form cysts *in vivo,* we next generated ΔCAP parasites and complemented versions in the type II Pru strain. In stark contrast to the RH line, upon injection of ~50,000 tachyzoites, Pru ΔCAP parasites showed no virulence in mice while the Pru ΔCAP^CAP^, Pru ΔCAP^shortCAP^, and Pru ΔCAP^longCAP^ complemented parasites led to a lethal infection, with most mice succumbing to the parasites 8-10 days post-infection (Fig. 8B). These data show that expression of either individual isoform is sufficient to cause a lethal infection. Next, to look at formation of tissue cysts, which leads to a chronic infection, we injected a lower dose of ~5,000 tachyzoites of the Pru lines into mice. The majority of mice survived until day 32 post-infection, although 2 ΔCAP^CAP^-infected mice and 1 ΔCAP^shortCAP^-infected mouse died before reaching this end-point. At day 32, the mice were sacrificed, brain samples were collected and serum tested for anti*-Toxoplasma* antibodies, confirming that all mice were successfully infected with *Toxoplasma* (Fig. 8C). Both the Pru ΔCAP and Pru ΔCAP^longCAP^ infections demonstrated significantly lower cyst loads compared to Pru ΔCAP^CAP^ and Pru ΔCAP^shortCAP^ infected mice (Fig. 8D). These results show that CAP plays an essential role in the virulence of the type II Pru parasite strain, but not the type I RH strain. Moreover, while the short CAP isoform is able to fulfil all functions of TgCAP in cell culture and in the mouse model, the long isoform, despite its ability to complement most phenotypes to at least ~50% of WT complement levels, has a significant defect in establishing a chronic infection at the infectious dose used here.

**Figure 8:**
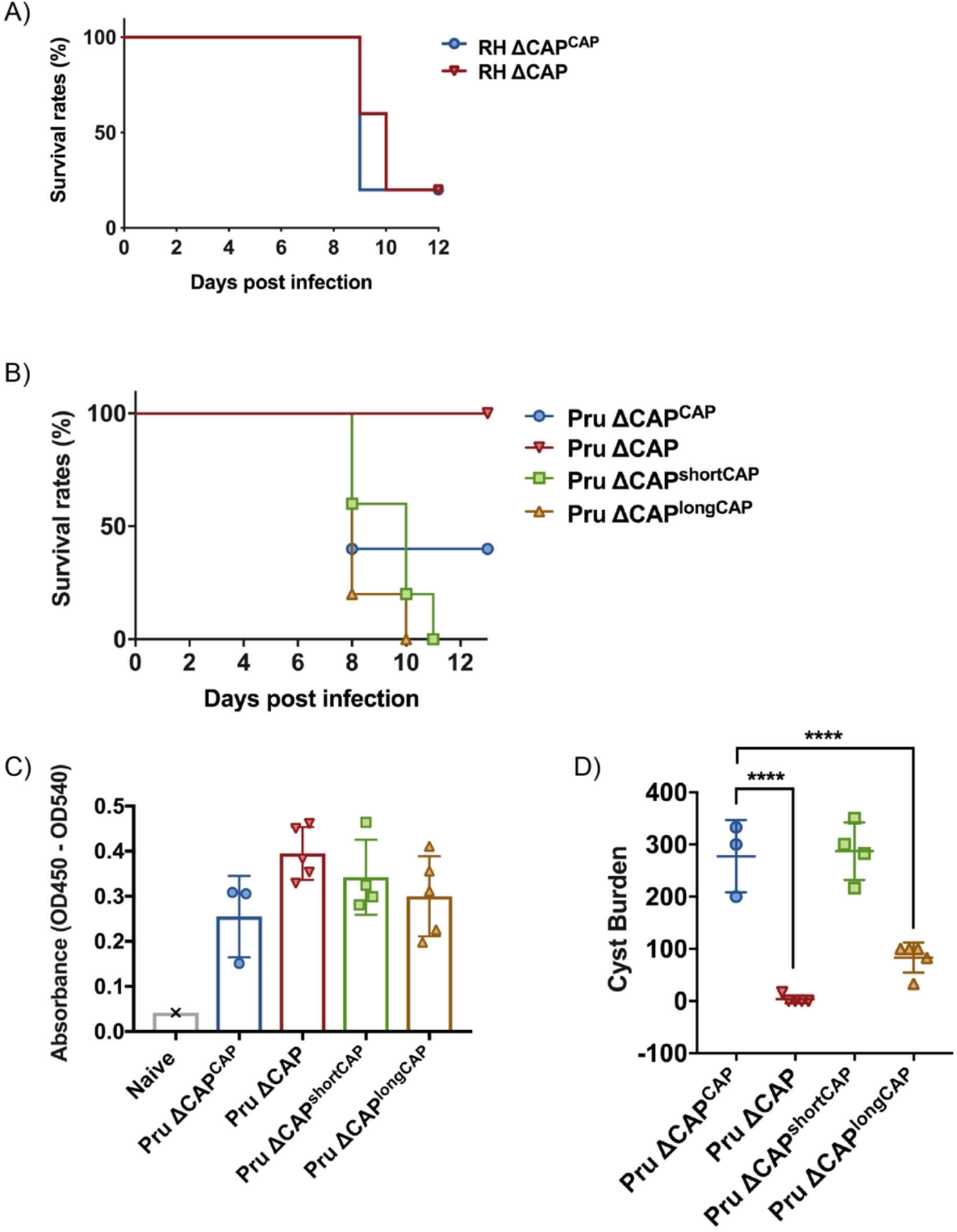
CAP is essential for virulence in type II but not type I parasites. **(A)** Survival rates of C57BL/6 mice infected with 25 RH ΔCAP^CAP^ or RH ΔCAP parasites. **(B)** Survival rates of C57BL/6 mice infected with 50,000 Pru ΔCAP parasites or with complementing CAP variants. **(C)** ELISA testing sera reactivity of naïve or *Toxoplasma* infected mice using *Toxoplasma* antigen. **D)** Cyst burden in the brain of C57BL/6 mice 32 days post-infection with 5,000 Pru ΔCAP parasites or with complementing CAP variants. For all experiments, five animals were infected for each strain.

## Discussion

*Toxoplasma* actin is important for a range of cellular processes, from organelle segregation and cell-cell communication, to gliding motility: a crucial factor in parasite dissemination (Andenmatten et al., 2013; Egarter et al., 2014; Periz et al., 2017). Despite the key role of actin in parasite biology, our understanding of actin dynamics and its regulation remains incomplete. CAP is a ubiquitous protein with a conserved role in regulating actin dynamics. In this study, we established that *Toxoplasma* CAP is expressed by an alternative translation initiation site, giving rise to two independent isoforms: shortCAP and longCAP. Through sequence alignment we identified that the alternate translation initiation site is conserved only within *Toxoplasma, Neospora and Hammondia.* These are all members of the Toxoplasmatinae, a subfamily of the Apicomplexa phylum, suggesting that while CAP is present in all apicomplexa, the long isoform is specific to *Toxoplasma* and its closest relatives. Here we investigated the role of both CAP isoforms in the *Toxoplasma* lytic cycle. Through *in vitro* competition assays, we show that complementation with shortCAP is enough to overcome the majority (87.6%), but not all, of the growth defect associated with CAP depletion. This suggests that longCAP is performing a specific function important for parasite fitness, which shortCAP is unable to compensate for. Furthermore, longCAP complementation restored 56.9% of the growth defect, arguing for a degree of functional overlap between the two isoforms. Our experiments have not uncovered a phenotype that only longCAP can rescue, which would have revealed a unique function. Its apical localisation, however, makes it temping to speculate a function in actin regulation during invasion, which is a phenotype it can fully rescue. It could be that higher actin turnover is required at the apex and the concentration of longCAP here is fulfilling this need by influencing G-actin levels. Indeed, it has previously been hypothesised in other organisms that increased local concentration of CAP results in the sequestration of actin monomers (Ono, 2013). However, providing evidence for this hypothesis is limited by the fact that the shortCAP isoform can fully rescue all phenotypes in cell culture under the conditions tested. Given that cell culture assays do not fully represent the environment *Toxoplasma* normally encounters, we aimed to tease apart the functions of the two isoforms in mice. Surprisingly, even here the short isoform appears able to compensate for the lack of longCAP. This suggests that even under conditions encountered in the natural host, the longCAP isoform plays only a minor role, although dosage effects or routes of infection may well be confounding factors when assessing virulence of the different strains. Competition experiments between the mutants *in vivo* may help to tease apart the independent importance of the isoforms in the future. However, from the *in vitro* competition assays it can be predicted that parasites expressing two isoforms fare better than those expressing just one isoform. We cannot rule out that the differences in proteins levels of longCAP and shortCAP in the single isoform producing lines are affecting the phenotypic observations. However, both isoforms are able to rescue most phenotypes indicating that the protein levels do not appear to substantially affect CAP function.

Despite being dispensable for type I RH parasite virulence, CAP is essential for type II Pru parasite virulence in mice, and complementation with either shortCAP or longCAP restored lethality. At lower, non-lethal doses, longCAP complementation led to markedly reduced cyst formation in the brain. The underlying basis for this has not been explored here and it could be that it is the reduced fitness of the strain, rather than stage conversion phenotype, that causes a reduction of parasites reaching the brain. Therefore, the observed virulence of the longCAP complemented strain is possibly due to the high dosage used, allowing the parasite to proliferate at high enough rates to overwhelm the immune system. Nevertheless, the basis for reduced fitness of ΔCAP and ΔCAP^longCAP^ parasites in the murine infection model is likely multifactorial and we hypothesise that other actin-dependent processes that we have not assayed here could also contribute to the reduced virulence in mice.

The most pronounced phenotype of CAP deletion is loss of rosetting: the highly symmetrical physical distribution of parasites within the vacuole. Previous studies on actin, myosin I, myosin J, ADF and formin 3 have suggested that the actomyosin motor is important for rosetting (Frénal, Dubremetz, et al., 2017; Haase et al., 2015; Periz et al., 2017; Tosetti et al., 2019). Our ΔCAP data further supports this hypothesis. Interestingly, complementing ΔCAP parasites with longCAP restored rosetting in just under half of all vacuoles, with the remainder appearing as equally disordered as the ΔCAP vacuoles. Despite this mixed population, we did not observe vacuoles with a combination of organised and disorganised parasites, suggesting rosetting is a binary outcome; either all parasites in a vacuole are connected by a central residual body, or not. This may suggest that the ability to produce highly organised rosettes is established during the first round of cell division when the residual body forms. This mixed population of rosetting and non-rosetting parasites, in an isogenic strain, gave us a unique opportunity to determine whether rosetting is important for efficient intravacuolar cell-cell communication. Using longCAP complemented parasites, we show that in rosetting vacuoles there is efficient RTORP with all parasites able to transfer material to the bleached parasite. Within the same population, non-rosetting vacuoles displayed severe defects in the RTORP with parasites seemingly only communicating in pairs. However, despite apparently only communicating in pairs, all parasites in the vacuole remained synchronised in their stage of replication. This is different to other studies which suggested that defects in RTORP between parasites leads to asynchrony in division (Frénal, Jacot, et al., 2017; Periz et al., 2017). Our results show that neither RTORP between daughter cells nor the formation of a rosette are predictors for synchronicity of division.

The synchronous division in ΔCAP parasites could be explained by our FIB SEM results. Despite their disorganised appearance and loss of rapid cell-cell communication, ΔCAP parasites are still connected by a decentralised residual body. This connection, although not facilitating rapid transfer of proteins between parasites that are further apart, could allow for slow or minimal transfer of proteins which is sufficient to synchronise divisions. An alternative hypothesis could be that metabolites, not proteins, are required to synchronise divisions, and their diffusion through the decentralised residual body is quicker. It is also a possibility that the seemingly continuous ER in the residual body, observed here connecting most if not all parasites, could contribute to synchronicity of division. Whatever the basis for synchronicity is, rosetting and rapid cell-cell communication are not essential and their analysis cannot be reliably used to predict whether connections between parasites exist. Recently, Foe *et al* reported that upon deletion of *ASH4,* a serine hydrolase, parasites were only connected in pairs but retained synchronicity of division (Foe et al., 2018), similar to our results. An explanation could be that ASH4 mutants display a phenotype similar to the CAP phenotype described here, where, despite the loss of RTORP, parasites remain connected by a decentralised residual body, allowing synchronicity of division.

In summary, our results strongly support an actin regulatory role for CAP in *Toxoplasma*. Interestingly, actin dependent processes were affected to differing extents in ΔCAP parasites, such as rosetting being completely lost while apicoplast inheritance was unaffected. This surely reflects the different spatial requirements for actin turnover within a cell. It is likely that the local concentration of actin, actin binding proteins such as the formins and the different myosins facilitate this. The results obtained here also leave open a few questions that are interesting to study in the future. How does actin help to initiate and maintain the centralised residual body? How are ER connections between parasites in a decentralised residual body maintained and what is the function of this connection? What is the functional role of longCAP with its high concentration at the apex of the parasite? Why are some actin dependent processes completely reliant on CAP while others are not? Each of these questions will require careful analysis and the cell lines described here will likely provide useful tools to investigate these in the future.

## Materials and methods

### Plasmid construction

All primers used in this study are listed in Supplementary File 1. All synthetic DNA used in this study is listed in Supplementary File 2.

To generate the CAP-GFP fusion plasmid, pUPRT_CAP_GFP, the *Toxoplasma CAP* gene (TGME49_310030) 5’UTR was amplified from genomic DNA using primer pair P1/P2 and Gibson assembled (Gibson et al., 2010) with a synthetic *CAP* cDNA-XmaI-eGFP sequence (GeneArt strings, Life Technologies, Massachusetts, United States) into *BamHI* and *PacI* digested UPRT targeting vector pUPRT-HA (Reese et al., 2011).

To generate the CAP cKO plasmid, pG140_CAP_cKO_LoxP111, the *CAP* 5’UTR with a LoxP site inserted 111 bp upstream of the *CAP* start codon, and a recodonised *CAP* cDNA-HA sequence, were synthesised (GeneArt strings, Life Technologies). These DNA fragments were Gibson cloned (Gibson et al., 2010) into the parental vector p5RT70loxPKillerRedloxPYFP-HX (Andenmatten et al., 2013) which had been digested using *ApaI* and *PacI,* creating an intermediate plasmid. Next, the *CAP* 3’UTR was amplified from genomic DNA using primer pair P3/P4 while mCherry flanked by *GRA* gene UTRs was amplified from pTKO2c (Caffaro et al., 2013) using primers P5/P6. These PCR products were Gibson cloned (Gibson et al., 2010) into the SacI digested intermediate plasmid to create pG140_CAP_cKO_LoxP111.

To generate pUPRT_CAP, the *CAP* 5’UTR was amplified from genomic DNA using primer pair P1/P2. This DNA fragment and a synthetic *CAP* cDNA-*BamHI*-HA sequence (GeneArt strings, Life Technologies), were Gibson cloned (Gibson et al., 2010) into *BamHI* and *PacI* digested UPRT targeting vector pUPRT-HA (Reese et al., 2011).

To generate pUPRT_CAP_C6,8A, the *CAP* 5’UTR was amplified from genomic DNA using primer pair P1/P2 and, alongside a synthetic *CAP* cDNA-HA sequence with C6,8A mutations (GeneArt strings, Life Technologies), was Gibson cloned (Gibson et al., 2010) into *BamHI* and *PacI* digested UPRT targeting vector pUPRT-HA (Reese et al., 2011).

To generate pUPRT_CAP_M1L, pUPRT_CAP was amplified with primer pair P7/P8 to introduce the M1L point mutation.

To generate pUPRT_CAP_M37L, pUPRT_CAP was amplified with primer pair P9/P10 to introduce the M37L point mutation.

To generate pG140_DiCre, the plasmid containing DiCre_T2A, two synthetic DNA fragments were produced containing *Ku80* homology region-alpha-tubulin promoter-FRB-Cre60-T2A-chloramphenicol resistance cassette-T2A-FKBP-Cre59-*SAG1* 3’UTR-*Ku80* homology region (gBlock gene fragments, Integrated DNA Technologies, Iowa, United States). These DNA fragments were Gibson cloned (Gibson et al., 2010) (into *ApaI* and *SacI* digested parental vector, p5RT70loxPKillerRedloxpYFP-HX (Andenmatten et al., 2013)

To generate plasmids expressing CAS9 and a single guide RNA (sgRNA), we used plasmid pSAG1::CAS9-U6::sgUPRT as a backbone (Addgene plasmid #54467) (Shen et al., 2014). The plasmid was amplified with primer P11 and another containing a sgRNA to replace the UPRT-targeting sgRNA. Guide RNA sequences were selected using the Eukaryotic Pathogen CRISPR gRNA Design Tool (Peng & Tarleton, 2015). All sgRNA-expressing plasmids used in this study were generated by this strategy. The sgRNA-containing primers are listed in Supplementary File 1.

For creation of plasmids with multiple sgRNAs, first, two separate vectors, each with a sgRNA, were generated as described above using the parental vector pSAG1::CAS9-U6::sgUPRT. Then, primers P12/P13 were used to amplify one of the sgRNA regions which was Gibson cloned (Gibson et al., 2010) into the other *KpnI* and *XhoI* digested plasmid, creating a multiple sgRNA plasmid.

### Culturing of parasites and host cells

*T. gondii* tachyzoites were cultured in human foreskin fibroblasts (HFF) and in Dulbecco’s modified Eagle’s medium (DMEM) with GlutaMAX (Invitrogen, California, United States) supplemented with 10% fetal bovine serum and maintained at 37°C with 5% CO_2_.

### Generation of parasite lines

To generate the conditional *CAP* knockout strain (RH DiCreΔ*ku80*Δ*hxgprt*_*LoxCAP-HA*, referred to here as LoxPCAP), first, the linearised plasmid pG140_CAP_cKO_LoxP111, carrying the *HXGPRT* cassette, was transfected into the RH DiCreΔ*ku80*Δ*hxgprt* strain (Andenmatten et al., 2013). Resistant parasites were cloned. Next, the DiCre conditional knockout function of the strain was restored. The linearised pG140_DiCre plasmid, carrying the chloramphenicol resistance cassette and homology with the *Ku80* UTRs, was transfected into the strain. To increase *Ku80*-specific insertion efficiency, a plasmid with multiple *Ku80*-targeting sgRNAs was generated as described above using primer pairs P11/P14 and P11/P15. Resistant parasites were cloned. Integration into the *CAP* endogenous locus was confirmed using primer pairs P16/P17 and P18/P19. Replacement of *CAP* gDNA was confirmed using primers P20/P21. Excision of the floxed *CAP* sequence was confirmed with primer pairs P16/P20.

To complement the LoxPCAP strain with *CAP*-expressing constructs, the linearised plasmid pUPRT_CAP, pUPRT_CAP_M1L or pUPRT_CAP_M37L was transfected alongside pSAG1::CAS9-U6::sgUPRT. 5-fluorodeoxyuridine (FUDR) resistant parasites were cloned.

*CAP* was subsequently excised from the above strains by addition of 50 nM rapamycin in DMSO for 4 hr at 37°C with 5% CO_2_, before washout, and excised parasites were cloned. Next, to aid with experimentation, an mCherry fluorescent construct was integrated into the *Ku80* locus, replacing the present DiCre_T2A construct. mCherry flanked by *GRA* UTRs was amplified from pG140_CAP_cKO_LoxP111 using primer pair P22/P23 which also carries 30 bp homology with the *Ku80* locus. To increase *Ku80*-specific insertion efficiency, a plasmid with multiple *Ku80*-targeting sgRNAs was generated as described above using primer pairs P11/P24 and P11/P25 and co-transfected with the PCR product. A population of parasites expressing mCherry were sorted by flow cytometry using a BD Influx cell sorter (BD Biosciences). Parasites were subsequently cloned, generating the strains ΔCAP, ΔCAP^CAP^, ΔCAP^shortCAP^ and ΔCAP^longCAP^.

The Pru Δ*ku80*Δ*cap* strain was generated by amplifying the *HXGPRT* resistance cassette from the pG140_CAP_cKO_LoxP111 plasmid using primer pair P26/P27. To direct insertion of the PCR product to the *CAP* locus, a plasmid with multiple *CAP*-targeting sgRNAs was generated as described above using primer pairs P11/P28 and P11/P29 and co-transfected with the PCR product. Resistant parasites were cloned. To complement the Pru Δ*ku80*Δ*cap* strain with *CAP*-expressing constructs, the linearised plasmid pUPRT_CAP, pUPRT_CAP_M1L or pUPRT_CAP_M37L was transfected alongside pSAG1::CAS9-U6::sgUPRT. FUDR resistant parasites were cloned.

To generate fluorescent Δ*gra2* parasites for FRAP experimentation, mCherry flanked by *GRA* UTRs was amplified from pG140_CAP_cKO_LoxP111 using primer pair P30/P31 with *UPRT* locus overhangs. This PCR product was co-transfected with pSAG1::CAS9-U6::sgUPRT into the RH Δ*ku80* Δ*gra2* strain (Rommereim et al., 2016). An FUDR resistant population was obtained.

The RH DiCre_T2A Δ*ku80*Δ*hxgprt* line was generated by integrating the DiCre construct into the *Ku80* locus in RH Δ*ku80*Δ*hxgprt* parasites (Huynh & Carruthers, 2009). The DiCre construct was amplified from pG140_DiCre using primer pair P32/P33 which also carries 30 bp homology with the *Ku80* locus. To increase *Ku80*-specific insertion efficiency, a plasmid with a *Ku80*-targeting sgRNA was generated as described above using primer pairs P34/P11 and co-transfected with the PCR product. Resistant parasites were cloned.

To assess the efficiency of rapamycin-dependent excision of the DiCre strains, the Killer Red gene-swap construct was amplified from p5RT70loxPKillerRedloxPYFP-HX (Andenmatten et al., 2013) using primer pair P35/P36. This PCR product was co-transfected with pSAG1::CAS9-U6::sgUPRT for targeted insertion into the *UPRT* locus. FUDR resistant parasites were cloned.

To generate a CAP C-terminal endogenous HA-tagged line, a synthetic DNA repair template was produced containing *CAP* gDNA (a section of which is recodonised), a HA tag and the *CAP* 3’UTR (gBlock gene fragments, Integrated DNA Technologies). The DNA repair template was amplified using primer pair P37/P38. To direct insertion of the PCR product to the *CAP* locus, a plasmid with a single *CAP*-targeting sgRNA was generated as described above using the primer pairs P39/P11 and was co-transfected with the PCR product into RH Δ*ku80* parasites. A population of parasites expressing the co-transfected Cas9-GFP-containing plasmid were sorted by flow cytometry using a BD Influx cell sorter (BD Biosciences). ~70% of these parasites expressed the HA peptide by IFA.

### Parasite transfection and selection

To generate stable transformants, 0.5 – 1 x10^7^ freshly lysed parasites were transfected with either 25 μg of linearised template DNA, 5 μg of linearised template DNA and 20 μg of a corresponding gRNA-specific CRISPR/CAS9 plasmid or template DNA produced from 1 ethanol precipitated PCR and 20 μg of a corresponding gRNA-specific CRISPR/CAS9 plasmid. Selection on the basis of 5-fluorodeoxyuridine (20 μM), mycophenolic acid (25 μg ml^-1^), xanthine (50 μg ml^-1^) or chloramphenicol (21 μM) was performed according to the selection cassette used.

### Preparation of parasite genomic DNA

Genomic DNA was extracted from *T. gondii* tachyzoites to use as a PCR template by pelleting parasites and resuspending in PBS. DNA extraction was then performed using the Qiagen QIAamp DNA blood mini kit as per the manufacturer’s protocol.

### IFA

Parasites were seeded onto HFFs grown on coverslips. 16 – 24 h after seeding, the coverslips were fixed in 3% formaldehyde for 15 min at room temperature then permeabilised in 0.2% Triton X-100/PBS for 3-10 min and blocked in 3% BSA/PBS for 1 h. Staining was performed using appropriate primary antibodies and goat Alexa Fluor 488-, Alexa Fluor 594- and Alexa Fluor 647-conjugated secondary antibodies (1:2000) alongside DAPI (5 μg/ml). Coverslips were mounted on glass slides with SlowFade gold antifade mountant (Life Technologies). Antibody concentrations used were: rat anti-HA high affinity (Roche, Basel, Switzerland) (1:1000), rabbit anti-TgCAP (1:2000), mouse anti-GFP (Roche) (1:1000).

Widefield images were generated with a Ti-E Nikon microscope using a 63x or 100x objective (Tokyo, Japan). Images were processed with Nikon Elements software. Confocal images were taken using a Zeiss LSM-780 inverted confocal laser scanning microscope with a 63x objective. Images were processed with Zeiss Zen Black software (Oberkochen, Germany).

### Western blot

Western blot samples were obtained by scraping and lysing intracellular parasites in 200 μl 1x Laemmli buffer (2% SDS, 10% glycerol, 5% 2-mercaptoethanol, 0.002% bromophenol blue and 125 mM Tris HCl, pH 6.8). Samples were subjected to SDS-PAGE under reducing conditions before being transferred to a nitrocellulose membrane. Immunoblotting was performed with appropriate primary antibodies in 0.1% Tween 20, 3% skimmed milk/PBS. Bound secondary fluorochrome-conjugated antibodies were visualised using the Odyssey Infrared Imaging System (LI-COR Biosciences, Nebraska, United States).

Antibody concentrations used were: rat anti-HA high affinity (Roche) (1:1000), rabbit anti-TgCAP (1:2000), mouse anti*-Toxoplasma* [TP3] (Abcam, Cambridge, United Kingdom) (1:1000). Goat anti-mouse IRDye 800CW (LI-cOr) (1:20000), Donkey anti-rabbit IRDye 680LT (LI-COR) (1:20000), Goat anti-rat IRDye 680LT (LI-COR) (1:20000).

### Generation of TgCAP antibody

To generate the shortCAP expression plasmid, pET-28_CAP_A38toC203, a *Toxoplasma* shortCAP (M37toC203) recodonised sequence was synthesised (gBlock gene fragments, Integrated DNA Technologies) and Gibson cloned (Gibson et al., 2010) into *BamHI* and *NdeI* digested pET-28a(+) plasmid (Merck, Darmstadt, Germany). This allowed for expression of an N-terminal 6xHis tagged shortCAP recombinant protein in *Escherichia coli* BL21 cells under the control of T7 *lac* promoter. Short CAP was expressed and His tag purified using Ni-NTA affinity purification under native conditions using the standard manufacturer’s protocol (Qiagen, Hilden, Germany). The shortCAP recombinant protein was used to immunize female New Zealand white rabbits (Covalab, Cambridge, United Kingdom) for generation of polyclonal antibodies.

### Flow cytometry analysis of DiCre excision

Parasites were added to a HFF monolayer and allowed to invade for 1 h. Then, cre recombinase-mediated recombination was induced by addition of 50 nM rapamycin in DMSO for 4 hr before washout. 22 h after infection, parasites were lysed, pelleted and washed twice in PBS. Parasites were resuspended in 0.5 ml 3% formaldehyde and fixed for 10 min. The suspension was centrifuged and the pellet washed in PBS before resuspension in PBS. To remove debris, samples were passed through a 30 μm pre-separation filter (Miltenyi Biotec, Bergisch Gladbach, Germany). 20,000 events were recorded using a BD LSR II flow cytometer (BD Biosciences California, United States). Killer Red was excited by the 561 nm laser and detected by a 600 long pass filter and either a 582/15, 610/20 or 620/40 band pass filter. YFP was excited by the 488 nm laser and detected a 505 long pass filter and either a 525/50 or 530/30 band pass filter. For quantification, and elimination of debris, total number of Killer Red+ parasites was considered 100%. This was performed at day 0, 35 and 65 of the experiment. For each condition, three biological replicates were analysed. At least 10000 Killer Red+ events were counted for each individual time point.

### Phenotypic characterisations

#### Competition Assay

mCherry-expressing ΔCAP, ΔCAP^CAP^, ΔCAP^shortCAP^ or ΔCAP^longCAP^ parasites were mixed with non-fluorescent ΔCAP^CAP^ parasites at an average ratio of 60/40. At day 0, 15 and 30 in culture, the ratio was determined by flow cytometry for two biological replicates. Parasites were lysed, pelleted and washed twice in PBS. Parasites were resuspended in 0.5 ml 3% formaldehyde and fixed for 5 min. The suspension was centrifuged and the pellet resuspended in 5 μg/ml DAPI/PBS for 10 min. The pellet was washed and resuspended in PBS. To remove debris, samples were passed through a 30 μm pre-separation filter (Miltenyi Biotec). Events were recorded using a BD LSR II flow cytometer (BD Biosciences). DAPI was excited by the 355 nm laser and detected by a 450/50 band pass filter. mCherry was excited by the 561 nm laser and detected by a 600 long pass filter and a 610/20 band pass filter. To eliminate debris from the analysis, events were gated on DAPI fluorescence. The ratio of control parasites (DAPI+/mCherry-) to individual CAP complements (DAPI+/mCherry+) was calculated and normalised to the day 0 ratio. The data represent two (day 15) and three (day 30) independent experiments. At least 1500 DAPI+ events were obtained for each individual time point. The results were statistically tested with a two-way ANOVA test plus a multiple comparison Sidak’s test individually comparing day 15 or day 30 means to their respective day 0 mean, in GraphPad Prism^®^ 7. The data presented are as mean ± s.d.

#### Intracellular growth and rosetting assay

Parasites were harvested from a T-25 and added to a coverslip coated with a HFF monolayer. After 20 h the coverslips were fixed with 3% formaldehyde for 15 min at room temperature. Coverslips were mounted and mCherry expression used to identify parasites. For each replicate, 4 random fields were imaged with a 40x objective. Counts were performed in a blinded manner in duplicate for two independent experiments. The number of parasites per vacuole was determined by counting at least 265 vacuoles per strain. The number of vacuoles that rosette was determined by looking at 8-pac vacuoles, at least 90 vacuoles per strain were counted. The results were statistically tested with a one-way ANOVA test plus a multiple comparison Dunnett’s test comparing all means to the ΔCAP^CAP^ mean in GraphPad Prism^®^ 7. The data presented are as mean ± s.d.

#### Invasion assay

Red/green invasion assays were performed. mCherry-expressing parasites were lysed in an invasion non-permissive buffer, Endo buffer (44.7 mM K_2_SO_4_, 10 mM MgSO_4_, 106 mM sucrose, 5 mM glucose, 20 mM Tris-H_2_SO_4_, 3.5 mg/ml BSA, pH 8.2). 250 μl of 8×10^5^ parasites/ml in Endo buffer were added to each well of a 24-well flat-bottom plate [Falcon], which contains a coverslip with a confluent HFF monolayer. The plates were spun at 129 x g for 1 min at 37°C to deposit parasites onto the monolayer. The Endo buffer was gently removed and replaced with invasion permissive medium (1% FBS/DMEM). These parasites were allowed to invade for 1 min at 37°C after which the monolayer was gently washed twice with PBS and fixed with 3% formaldehyde for 15 min at room temperature. Extracellular parasites were stained with mouse anti*-Toxoplasma* antigen B1247M (Abcam) 1:1000 and goat anti-mouse Alexa Fluor 488, following the IFA protocol. For each replicate, 3 random fields were imaged with a 40x objective. Three independent experiments were performed in duplicate. The number of intracellular (mCherry+/488-) and extracellular (mCherry+/488+) parasites was determined by counting, in a blinded fashion, at least 758 parasites per strain. The results were statistically tested with a one-way ANOVA test plus a multiple comparison Dunnett’s test comparing all means to the ΔCAP^CAP^ mean in GraphPad Prismg^®^ 7. The data presented are as mean ± s.d.

#### Egress assay

Parasites were added to a HFF monolayer, in a ibidi μ-plate 96 well, and grown for 30 h. The wells were washed twice with PBS and the media was exchanged for 80 μl Ringers solution (155 mM NaCl, 3 mM KCl, 2 mM CaCl_2_, 1 mM MgCl_2_, 3 mM NaH_1_PO_4_, 10 mM HEPES, 10 mM glucose). To artificially induce egress, 40 μl of Ringer’s solution containing 150 μM BIPPO (50 μM final conc) was added to each well. At specified time points the cells were fixed by adding 26 μl 16% formaldehyde (3% final conc) for 15 min. Cells were washed in PBS and stained with DAPI (5 μg/ml). Automated image acquisition of 25 fields per well was performed on a Cellomics Array Scan VTI HCS reader (Thermo Scientific, Massachusetts, United States) using a 20× objective. Image analysis was performed using the Compartmental Analysis BioApplication on HCS Studio (Thermo Scientific). Egress levels were determined in triplicate for three independent assays. At least 11987 vacuoles per strain were counted at t = 0 s. Subsequent time point vacuole counts were normalised to t=0 to determine how many vacuoles had egressed. The results were statistically tested with a two-way ANOVA test plus a multiple comparison Dunnett’s test comparing all means to the ΔCAP^CAP^ mean, at each time point separately, in GraphPad Prism^®^ 7. The data presented are as mean ± s.d.

#### Live egress

Parasites were added to a HFF monolayer, in a ibidi μ-plate 96 well, and grown for 30 h. The wells were washed twice with PBS and the media was exchanged for 80 μl Ringers solution (155 mM NaCl, 3 mM KCl, 2 mM CaCl_2_, 1 mM MgCl_2_, 3 mM NaH_2_PO_4_, 10 mM HEPES, 10 mM glucose). The plate was then transferred to a Ti-E Nikon microscope with a 37°C environmental chamber. To artificially induce egress, 40 μl of Ringer’s solution containing 150 μM BIPPO (50 μM final conc) was added to each well after imaging had commenced. Images were captured every 1.8 s.

#### Apicoplast segregation assay

Parasites were added to a HFF monolayer and grown for 20 h before fixation with ice cold methanol for 2 min at room temperature. IFAs were performed using a streptavidin-Alexa Fluor 594 conjugate (Invitrogen) as a marker for the apicoplast. Correct apicoplast segregation was determined in duplicate for three independent assays. At least 228 vacuoles were counted per strain. Counts were performed in a blinded manner. The results were statistically tested with an unpaired t test in GraphPad Prism^®^ 7. The data presented are as mean ± s.d.

#### Synchronicity of division and daughter cell orientation assay

Parasites were added to a HFF monolayer and grown for 20 h before fixation with 3% formaldehyde for 15 min at room temperature. IFAs were performed using rat anti-IMC3 antibodies. To determine the synchronicity of cell division within the vacuoles, anti-IMC3 staining was used to evaluate the stage of daughter cell development. Vacuoles were scored as synchronous if all daughter cells were at the same stage of development. Vacuoles were counted blind in triplicate for three independent experiments. At least 275 vacuoles were counted per strain. The results were statistically tested with an unpaired t test in GraphPad Prism^®^ 7. The data presented are as mean ± s.d. Daughter cell orientation was quantified in triplicate for three independent experiments. At least 312 mother cells were counted blind, per strain. The results were statistically tested with a one-way ANOVA test plus a multiple comparison Dunnett’s test comparing all means to the ΔCAP^CAP^ mean in GraphPad Prism^®^ 7. The data presented are as mean ± s.d.

#### 3D motility assay

Motility assays were performed as previously described (Leung et al., 2014), with minor modifications. Parasites were syringe-released from a HFF monolayer (one heavily infected T75 flask per strain) by passing through a 27 gauge needle and filtering through a 3 μm Nuclepore filter. Parasites were then centrifuged (1,000 x g for 2 min) and resuspended in 40 μl motility media supplemented with 0.3 mg/mL Hoescht 33342. Matrigel was thawed on ice to prevent polymerization and combined with parasites and motility media in a ratio of 3:1:3 respectively. Pitta chambers were perfused with 10 μL of this suspension and incubated at 27°C for 7 minutes on a thermoplate. Chambers were incubated in the heated (35°C +/− 1°C) microscope enclosure for 3 min prior to imaging. Parasite nuclei were imaged, using a 20x objective, capturing 61 – 63 stacks of 41 z-slices 1 μm apart. To ensure conditions remained constant between parasite lines, samples used for capture were alternated. Datasets were exported to Imaris x64 v9.2.1. Using the ImarisTrack module parasites were tracked in a region of interest from which 1 μm has been cropped from the x and y edges to eliminate edge artefacts. Parasites were identified, after background subtraction, using spot detection with estimated size of 4 μm (xy) and 8 μm (z). Spots were filtered to exclude all that had duration of less than 3 seconds to minimise tracking artifacts. An autoregressive motion tracking algorithm was applied with a maximum distance of 6 μm and a maximum gap size of 3. Datasets were manually inspected to ensure appropriate tracking and to remove artifacts and trajectories tracked from multiple identified spots. Percent moving was calculated as the number of trajectories (>2 μm displacement) / number of objects in frame 3 (>3 s duration). Trajectory parameters were extracted directly from Imaris software. Data shown are derived from 4 independent biological replicates, each consisting of a minimum of two technical replicates. For all analysis, means and standard deviation were calculated for four independent biological replicates before statistical analysis using an unpaired t-test in GraphPad Prism ^®^7. For the three strain experiment, ΔCAP^shortCAP^ and ΔCAp^longCAP^ were each compared to ΔCAP^cap^. All technical replicates are presented in the figures, together with their mean ± s.d.

#### Dense granule trafficking assay

For each condition 1×10^7^ parasites were transiently transfected with 30 μg of pTub SAG1ΔGPI-mCherry plasmid (Heaslip et al., 2016) and immediately added to a confluent HFF monolayer in a Mattek^®^ 35 mm dish, coverslip 1.5. 12-15 h after infection the monolayers were washed 3 times with pre-warmed Gibco^®^ Fluorobrite™ DMEM supplemented with 4% Fetal Bovine Serum. The coverslips were immediately used for imaging. The acquisitions were made with an Olympus IX71 coupled to a DELTAVISION™ Elite imaging system in a 37°C environmental chamber. The acquisition for each condition were made sequentially from the washing step to the acquisition with a random order to avoid any artifactual data. The acquisition analysis was made with Fiji and the MTrackJ plugin. The data represent three independent experiments. At least 43 parasites and 273 direct runs were counted per strain. The results were statistically tested with a one-way ANOVA test plus a multiple comparison Dunnett’s test comparing all means to the ΔCAP^CAP^ mean in GraphPad Prism^®^ 7. The data presented are as mean ± s.d.

#### FRAP

Parasites were inoculated on a confluent layer of HFFs 20 h before experiments were performed using a Zeiss LSM-780 inverted confocal laser scanning microscope at 37°C. Acquisition and processing were performed with the Zeiss Zen Black software. Images were taken for 2 min (one image per second). Three pre-bleach images were recorded before the region of interest was photobleached ten times with a 561 nm laser at 100% power. Fluorescence intensity is presented as a percentage relative to the same area pre-bleach. These normalised intensity values were also used for the calculations below. To calculate percentage of fluorescence recovery, the final reading (116 s post-bleach) was subtracted from the reading immediately post-bleach, t = 3 s, to give percentage recovery after 116 s. To calculate percentage loss of fluorescence, the reading immediately post-bleach, t = 3 s, was subtracted from the final reading (116 s post-bleach) to give percentage loss after 116 s. These percentages were used to generate the heat map. At least 11 vacuoles were counted per strain across at least two independent experiments. The results were statistically tested with a one-way ANOVA test plus a multiple comparison Dunnett’s test comparing all means to the ΔCAP^CAP^ mean in GraphPad Prism^®^ 7. The data presented are as mean ± s.d. To assess recovery type, at least 11 vacuoles were counted per strain across at least two independent experiments. The results were statistically tested with a Chi-square in GraphPad Prism^®^ 7.

#### Tape unroofing SEM

Parasites were inoculated on a confluent HFF monolayer 24 h before fixation in EM fixative (2.5% gluteraldehyde, 4% formaldehyde in 0.1 M phosphate buffer) for 30 min. Cells were washed in 0.1 M phosphate buffer (PB) and stored in 1% formaldehyde in PB at 4°C. Cells were then washed in PB at room temperature, then washed in ddH_2_O at RT. The cells were dehydrated stepwise from 70% to 100% ethanol before critical point drying from acetone in a CPD300 (Leica Microsystems, Vienna, Austria). After drying, the coverslips were mounted on stubs, and the HFF cells were unroofed by placing Scotch tape on the coverslips and gently peeling it off, exposing the host cytoplasm and the parasitophorous vacuoles. The cells were coated with 7 nm platinum in a Q150R Sputter Coater (Quorum Tech, East Sussex, UK) before viewing in a Phenom ProX SEM (Thermo Scientific) at 10 kV, 1024 x 1024 pixel frame, on ‘high’ quality.

#### FIB SEM

Immediately following FRAP experimentation, as described above, parasites were fixed in 4% formaldehyde for 15 min at 37°C before washing in 0.1 M PB. Cells were then fixed in EM fixative (2.5% gluteraldehyde, 4% formaldehyde in 0.1 M PB) for 30 min at room temperature. Cells were washed in 0.1 M PB and stored in 1% formaldehyde in 0.1 M PB. After fixation, samples were transferred to a Pelco BioWave Pro+ microwave (Ted Pella) for processing using a protocol adapted from the NCMIR protocol (Deerinck et al., 2010). See Supplementary File 3 for full BioWave program details. The SteadyTemp plate was set to 21°C unless otherwise indicated. Each step was performed in the microwave, except for the buffer and ddH2O wash steps, which consisted of two washes on the bench and two washes in the microwave (250 W for 40 s). The cells were washed (as above) in 0.1 M PB, stained with 2% osmium tetroxide and 1.5% potassium ferricyanide (v/v) for 14 min under vacuum (with/without 100 W power at 2 min intervals), and then washed in ddH_2_O (as above). Next, the cells were incubated in 1% thiocarbohydrazide in ddH_2_O (w/v) for 14 min (vacuum, 100 W on/off at 2 min intervals) with SteadyTemp plate set to 40°C, followed by ddH_2_O washes (as above), and then a further stain with 2% osmium tetroxide in ddH_2_O (w/v) for 14 min (vacuum, 100 W on/off at 2 min intervals), followed by ddH_2_O washes (as above). The cells were then incubated in 1% aqueous uranyl acetate (vacuum, 100 W on/off at 2 min intervals, SteadyTemp 40°C), and then washed in ddH_2_O (as above, except with SteadyTemp at 40°C). Walton’s lead aspartate was then applied (vacuum, 100 W on/off at 2 min intervals, SteadyTemp 50°C), and the cells were washed (as above) and dehydrated in a graded ethanol series (70%, 90%, and 100%, twice each), at 250 W for 40 s without vacuum. Exchange into Durcupan ACM^®^ resin (Sigma-Aldrich, Missouri, United States) was performed in 50% resin in ethanol, at 250 W for 3 min, with vacuum cycling (on/off at 30 sec intervals), and then pure Durcupan was infiltrated in four microwave steps with the same settings, before embedding at 60°C for 48 h.

Focused ion beam scanning electron microscopy (FIB SEM) data was collected using a Crossbeam 540 FIB SEM with Atlas 5 for 3-dimensional tomography acquisition (Zeiss). Segments of the cell monolayer containing the cells of interest were trimmed, polished with a diamond knife (removing uneven resin at the base of the monolayer to provide a flat surface for tracking marks), mounted on a standard 12.7 mm SEM stub using conductive epoxy (ITW Chemtronics), and coated with a 5 nm layer of platinum.

The specific cells of interest were relocated by imaging through the platinum coating at an accelerating voltage of 20 kV and correlating to previously acquired fluorescence microscopy images. After preparation for milling and tracking, images were acquired at 5 nm isotropic resolution throughout each region of interest, using a 10 μs dwell time. During acquisition the SEM was operated at an accelerating voltage of 1.5 kV with 1 nA current. The EsB detector was used with a grid voltage of 1,200 V. Ion beam milling was performed at an accelerating voltage of 30 kV and current of 700 pA.

After cropping to the specific region of interest comprising the entire extent of the PV (ΔCAP^CAP^; 3080 x 634 x 2509 pixels; 15.4 x 3.17 x 12.545 μm; ΔCAP; 4065 x 1136 x 4490 pixels; 20.325 x 5.68 x 22.45 μm) and aligning the dataset (gradient align; Atlas 5), the images were processed to supress noise and slightly enhance sharpness (gaussian blur 0.75 radius, followed by unsharp mask radius 1, strength 0.6; Fiji) prior to reorienting and reslicing in the YZ plane (assigned as the XY plane for segmentation), and scaling to 10 nm isotropic resolution for segmentation and display.

Selected structures were segmented manually from the FIB SEM datasets and 3D reconstructions were made using the 3dmod program of IMOD (Kremer et al., 1996). The normal and decentralised residual body structures were manually skeletonised by tracing the approximate central axis of the structure from the posterior pore of each tachyzoite using open contours. Thus, the green skeleton follows the lumen of the connections between parasites.

The minimum number of points were placed that would still ensure the contour remained at the central axis of the volume. Where the structure branched, points linking contours from multiple extensions were placed at the approximate centre of the branch point volume. The contours were then rendered as a 50 or 100 nm tube to aid visualisation (tube diameter chosen depending on view). This model was then inspected in the X, Y, and Z-planes and corrections made to ensure the skeleton followed the approximate centre through the volume. Since not all RB-like structure extensions ended at the posterior pole of a tachyzoite, the posterior poles were highlighted by segmenting them with open contours with points at every 40 nm in Z, following the edge of the cytosol (the ribosome-containing electron lucent space where it meets the intermediate electron density surrounding the inner membrane complex), and meshing the contour as a 50 nm tube. A selected region of the putative ER lipid bilayer outer leaflet was segmented with closed contours drawn every 5-20 nm in Z (smaller Z intervals where needed to capture fenestrations/complexity); from an approximately square region around the inner face of two of the basal pores up to an arbitrary point along the decentralised residual body structure. Contour gaps were placed at the edge of this region. Two tachyzoites were also highlighted by partial coarse segmentation; closed contours drawn every 250 nm in Z through the main body of the cell (segmentation of the complex top and bottom of the tachyzoites was omitted for clarity).

#### Animals

C57BL/6 (wild type) mice were bred and housed under pathogen-free conditions in the biological research facility at the Francis Crick Institute in accordance with the Home Office UK Animals (Scientific Procedures) Act 1986. All work was approved by the UK Home Office (project license PDE274B7D), the Francis Crick Institute Ethical Review Panel, and conforms to European Union directive 2010/63/EU. All mice used in this study were male and between 7- to 9-week old.

Mice were infected with *T. gondii* tachyzoites by intraperitoneal injection (i.p.) with either 25, 5 x 10^3^ parasites (cyst formation) or 5 x 10^4^ parasites (survival) in 200 μl medium on day 0. Mice were monitored and weighed regularly for the duration of the experiments.

For serum samples, mice were euthanized and blood collected into blood serum collection tubes (SAI, Infusion technologies) by puncturing the jugular vein. Blood was allowed to clot at room temperature for 30 min, before tubes were centrifuged at 1500 x g for 10 min. Serum was collected and stored at −20°C until analysis.

#### *Toxoplasma* serum antibody ELISA

*Toxoplasma* soluble antigens were extracted as previously described (Silva et al., 2007). In short, parasites were syringe-lysed, washed once with PBS, and adjusted to 1 x 10^8^ tachyzoites/ml with PBS containing protease inhibitors (cOmplete mini, Roche). Parasites were lysed by five freeze-thaw cycles (liquid nitrogen/37°C), followed by ultrasound sonication on ice (five 60 Hz cycle for 1 min each). Samples were centrifuged at 10,000 x g for 30 min at 4 °C before supernatants were collected and protein content was determined using the BCA Protein Assay Kit (Pierce, Thermo Fisher Scientific) following the manufacturer’s instructions.

To detect *Toxoplasma* antibodies in murine serum samples, 96-well plates (flat bottom, high-binding) were coated overnight with 2 μg/ml *Toxoplasma* soluble antigens at 4°C. Plates were washed with PBS/0.05% Tween-20 (v/v) (PBS-T) before blocked with 1% BSA (w/v) in PBS for 2 h at room temperature. Bound antigens were incubated with murine sera diluted 1/10 in 1% BSA/PBS for 2 h at room temperature, washed three times with PBS-T and bound antibodies detected by incubation for 2 h at room temperature with anti-mouse Immunoglobulins (HRP conjugate, Darko) diluted 1/1000 in 1 % BSA/PBS. Finally, plates were washed three times with PBS-T and developed by adding TMB substrate solution (Thermo Fisher Scientific). The TMB reaction was stopped by adding 2 N sulphuric acid and the absorbance measured (OD_450_ minus OD_540_ wave length correction) using the VersaMax™ Microplate Reader with SoftMax^®^ Pro Software.

#### Mouse brain collection and preparation for cyst counting

To determine the number of cysts in the brain of infected animals, mice were euthanized and the brain extracted from the skull. The brain was homogenised in 1 ml PBS and stained with Rhodamine-conjugated *Dolichos biflorus* agglutinin (1/1000; Vector Laboratories) for 1 h at room temperature. Fluorescently labelled cysts were counted using a Ti-E Nikon microscope.

#### Statistical analysis

Statistical tests used are stated in individual sections above. *P*-values significance thresholds were set at: **** *P* <0.0001, *** *P* <0.001, ** *P* <0.01 and * *P* <0.05. All significant results are labelled with a line and asterisk(s) in the graphs.

## Acknowledgements

We thank Marc-Jan Gubbels (Boston College) for gifting the IMC3 antibody, Markus Meissner (University of Glasgow) for the pG140 plasmid, Michael Reese (University of Texas Southwestern Medical Center) for the pUPRT-HA plasmid, David Sibley (Washington University) for the CRISPR/Cas9 plasmid and Caia Dominicus (Francis Crick Institute) for critically reading the manuscript. We also thank the following members of science technology platforms at the Francis Crick Institute for their support: Matt Renshaw (light microscopy), Michael Howell (high throughput screening), Damian Carragher, Rhys Hefin, Phil Hobson and Graham Preece (flow cytometry). This work was supported by awards to MT by The Francis Crick Institute (https://www.crick.ac.uk/), which receives its core funding from Cancer Research UK (FC001189; https://www.cancerresearchuk.org), the UK Medical Research Council (FC001189; https://www.mrc.ac.uk/) and the Wellcome Trust (FC001189; https://wellcome.ac.uk/). This work was supported by US Public Health Service grants AI137767 and AI139201 to GEW and National Institutes of Health grant awarded to Aoife Heaslip (AI121885).

---

**Supplementary Figure 1.**
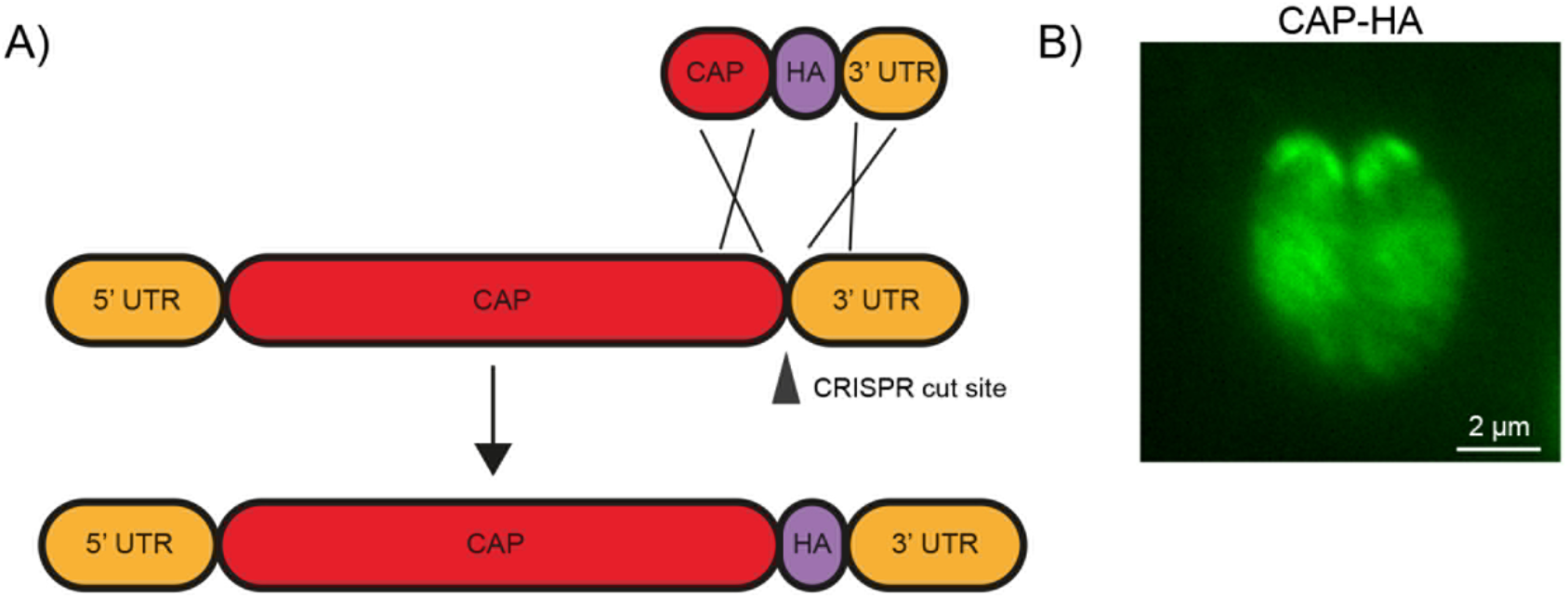
Endogenous tagging of CAP. **(A)** Schematic of CAP endogenous tagging strategy. Arrowhead indicates CRISPR/CAS9 cut site. **(B)** IFA of the resulting HA-tagged line. Scale bar, 2 μm.

**Supplementary Figure 2.**
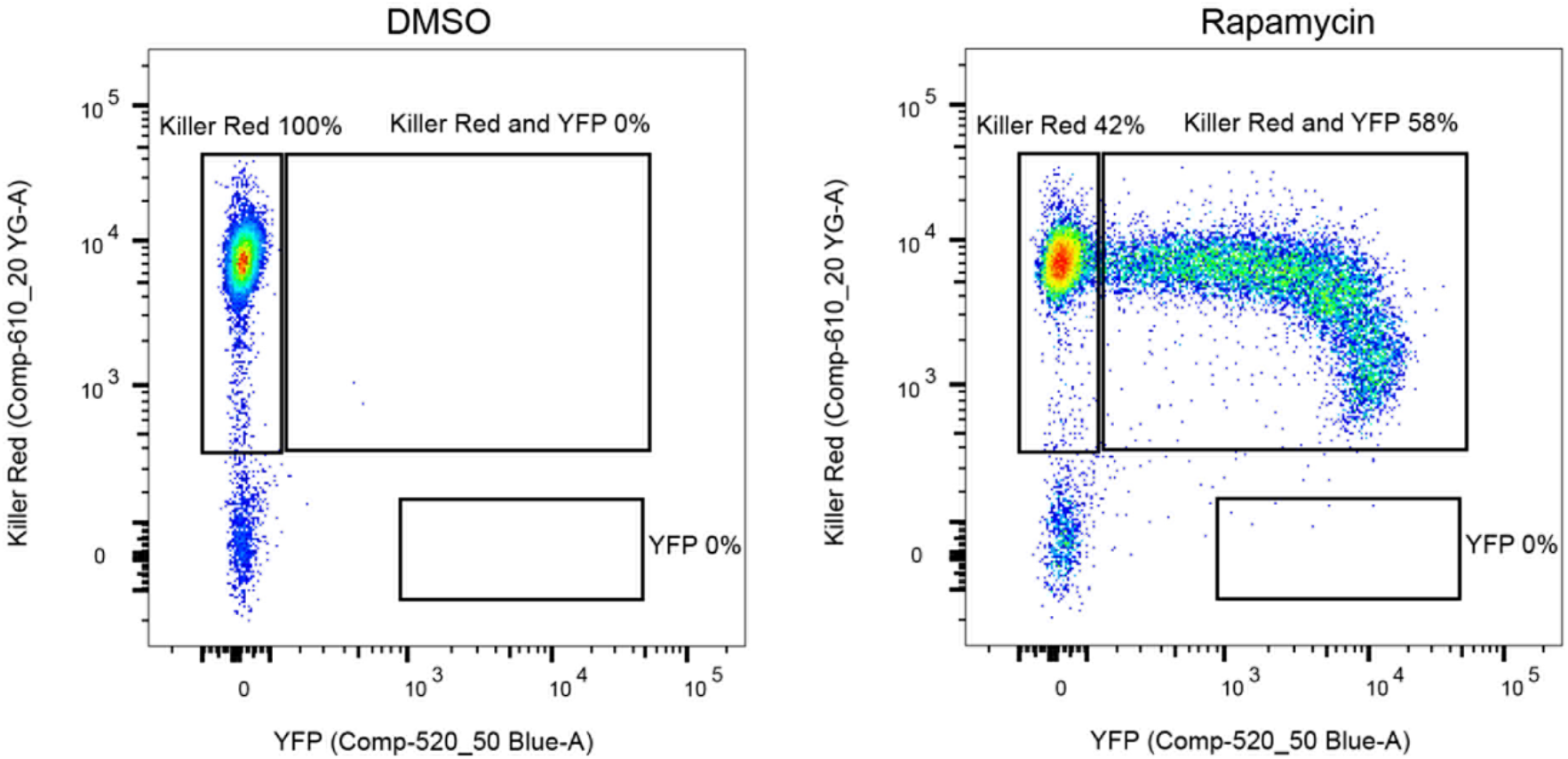
Excision efficiency testing of RH DiCre Δ*ku80*Δ*hxgprt*. Flow cytometry analysis to determine excision efficiency of the original RH DiCre Δ*ku80*Δ*hxgpr*t** line following 65 days of standard, non-stressing, culture conditions. Excision is determined by a shift from Killer Red^(+)^ to Killer Red^(+)^ and YFP^(+)^ expression. Parasites were analysed 22 h after induction with 50 nM rapamycin (RAP) for 4 h. Due to analysing 22 h after induction of excision, parasites still have residual KillerRed signal.

**Supplementary Figure 3.**
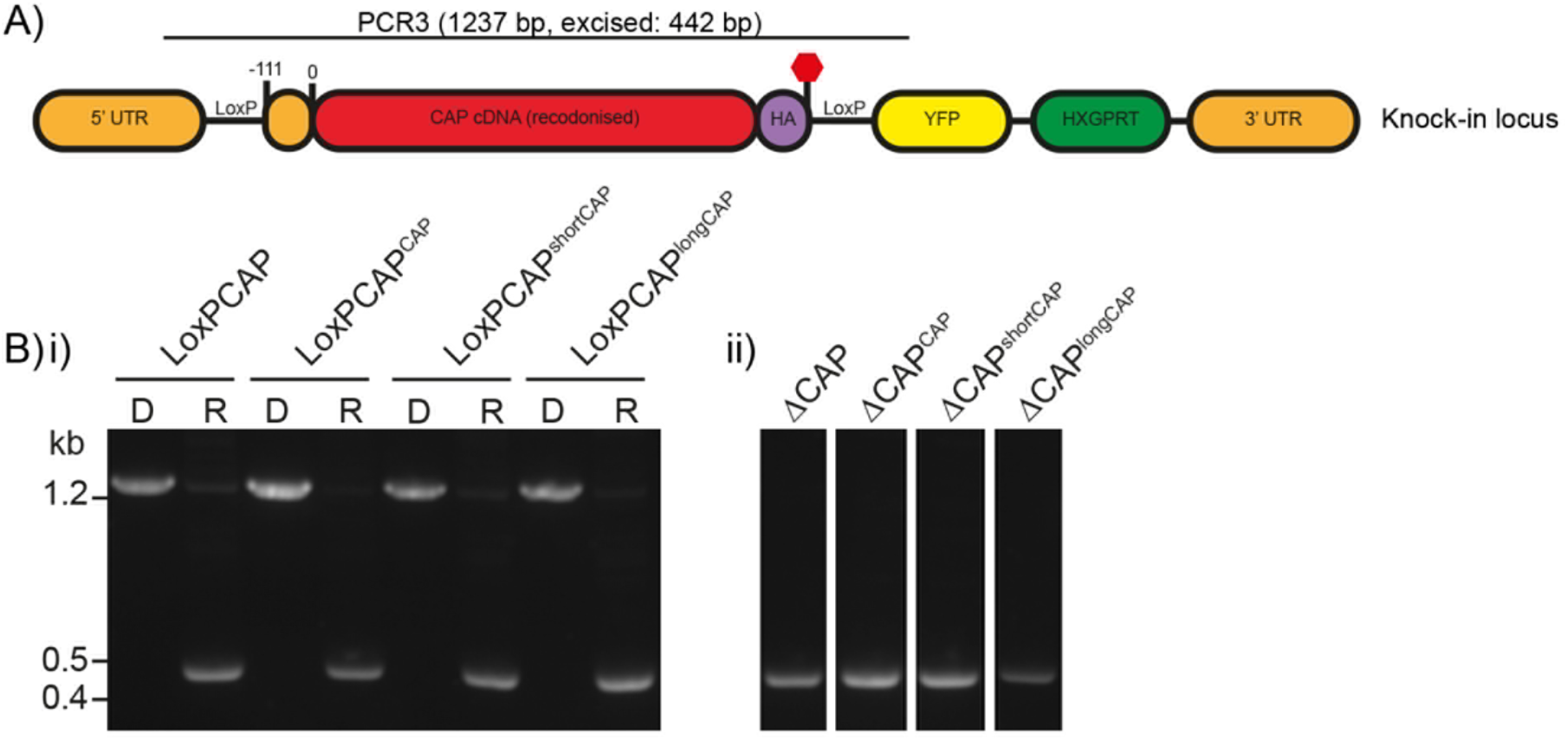
Excision testing of the CAP conditional knockout lines. **(A)** Schematic of the *CAP* locus in the LoxPCAP conditional knockout lines. Rapamycin induced excision of CAP is detected by PCR3. **(B)** Diagnostic PCR of the *CAP* locus in the **(i)** LoxPCAP lines 5 d after treatment with either DMSO (D) or rapamycin (R). **(ii)** Excised clones were subsequently obtained from these rapamycin treated populations.

**Supplementary Figure 4.**
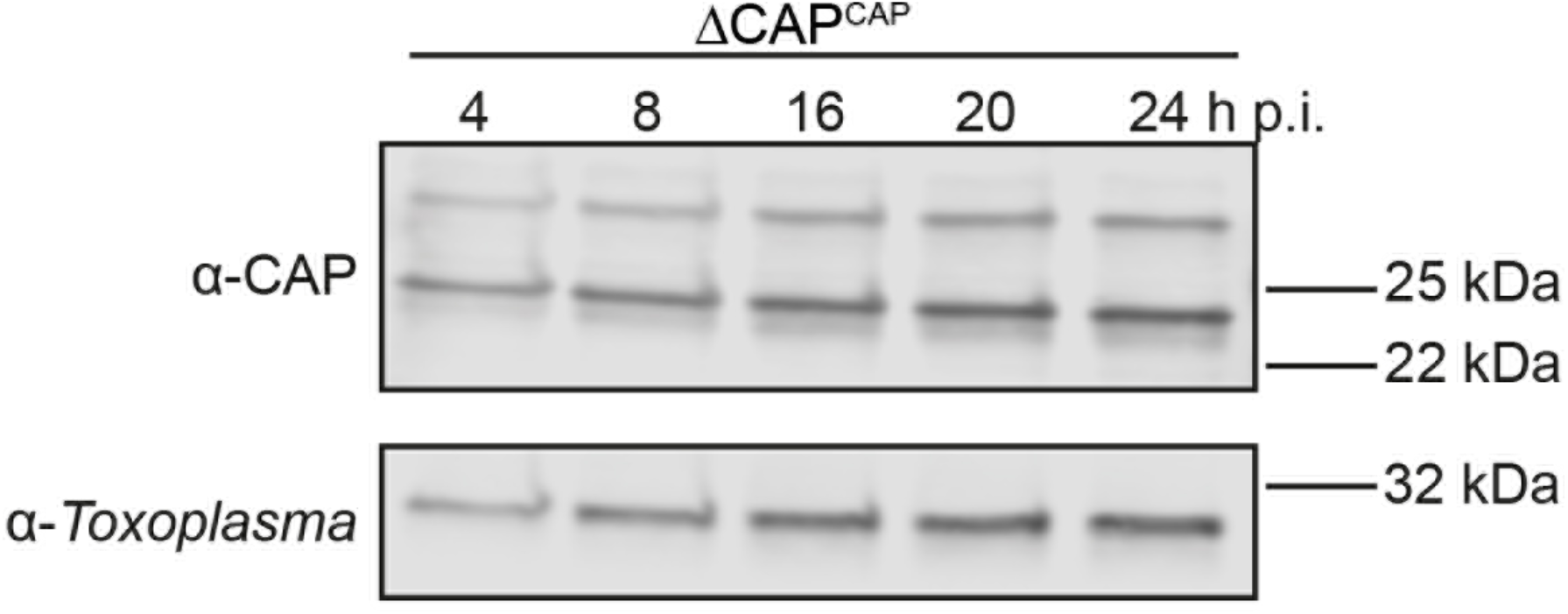
CAP-HA expression in the ΔCAP^CAP^ line. Western blot on CAP expression levels in the ΔCAP^CAP^ strain over the first 24 hours following host cell invasion using anti-TgCAP antibodies. Anti*-Toxoplasma* antibodies were used as a loading control.

**Supplementary Figure 5.**
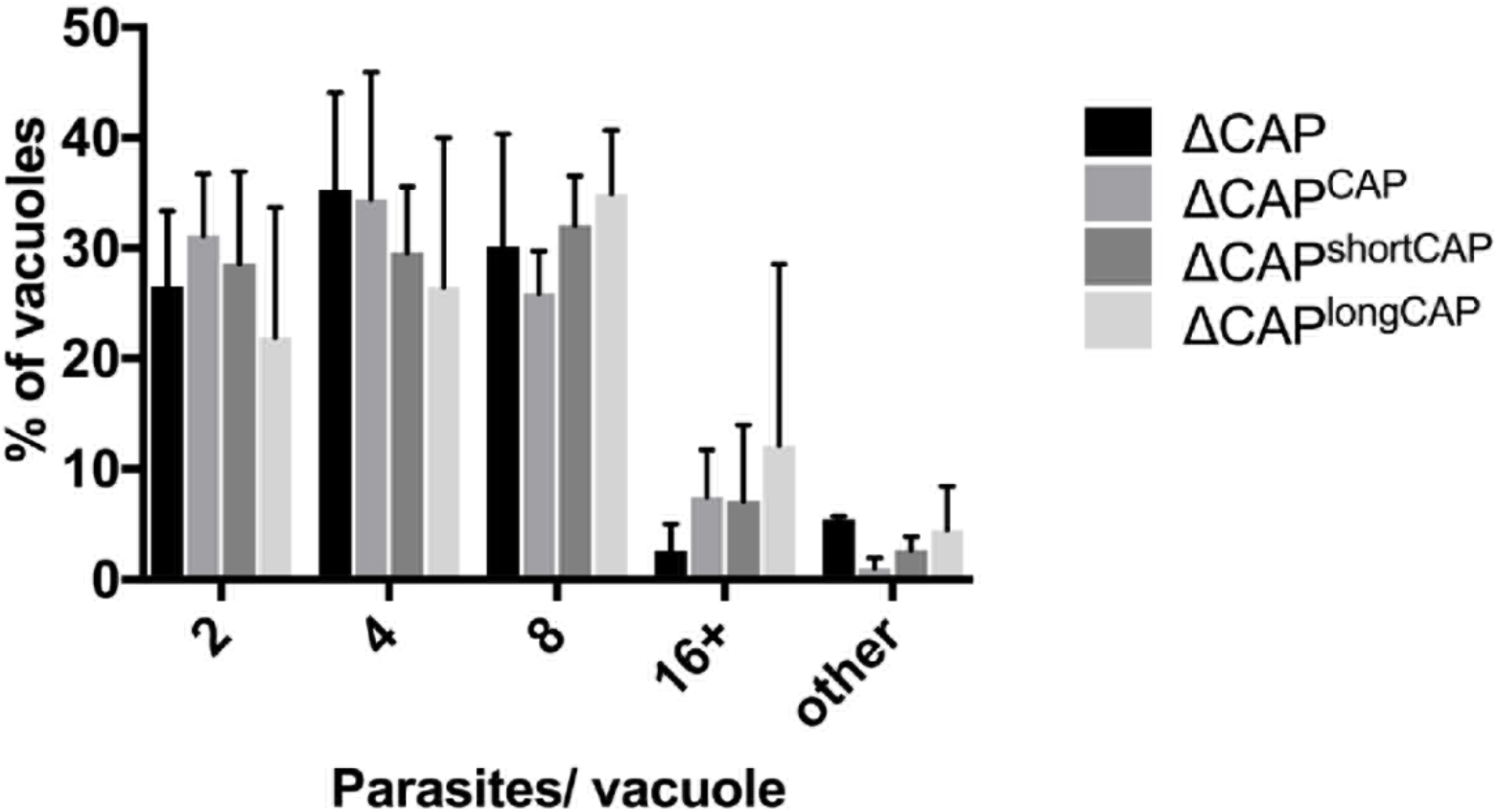
Replication analysis of ΔCAP and the complemented lines. Data are represented as mean ± s.d. (*n*=2). One-way ANOVA followed by Dunnett’s test was used to compare means to the ΔCAP^CAP^ mean. “Other” refers to vacuoles with a non-power of 2 number of parasites.

**Supplementary Figure 6.**
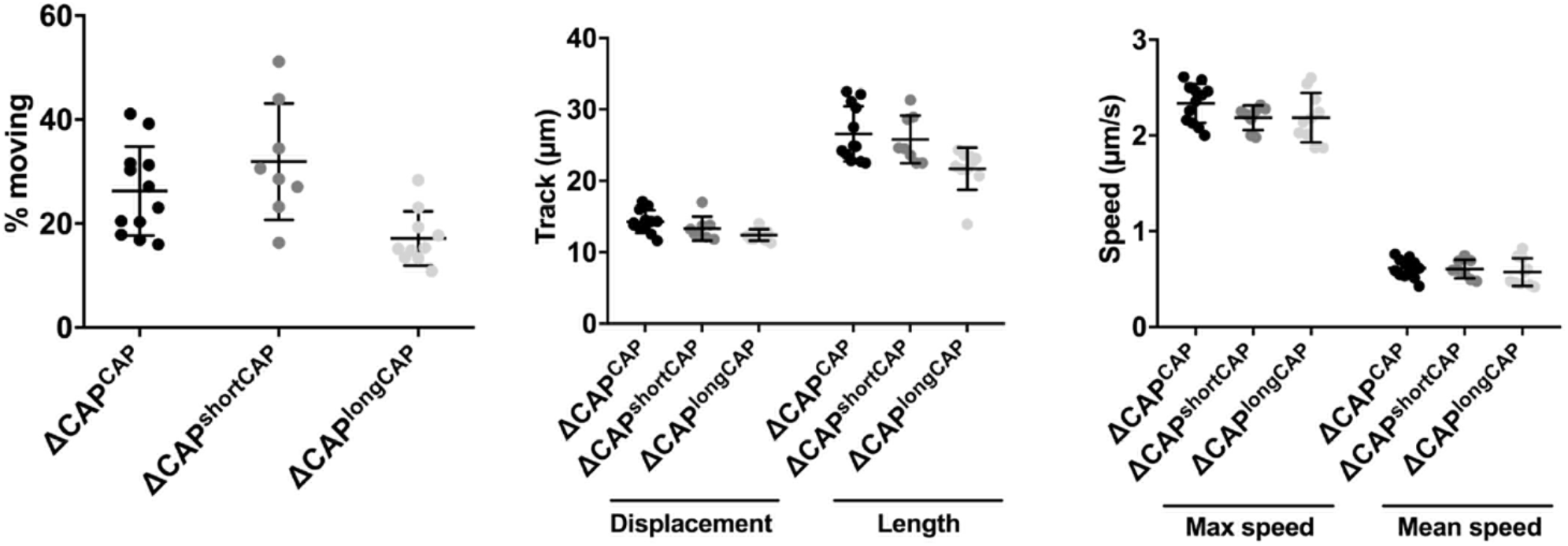
3D Matrigel-based motility assays performed in the absence of inducers of motility. Results are expressed as mean ± s.d. (*n*=4). Each data point corresponds to a single technical replicate from one of four independent biological replicates, on which significance was assessed using an unpaired *t*-test.

**Supplementary Figure 7.**
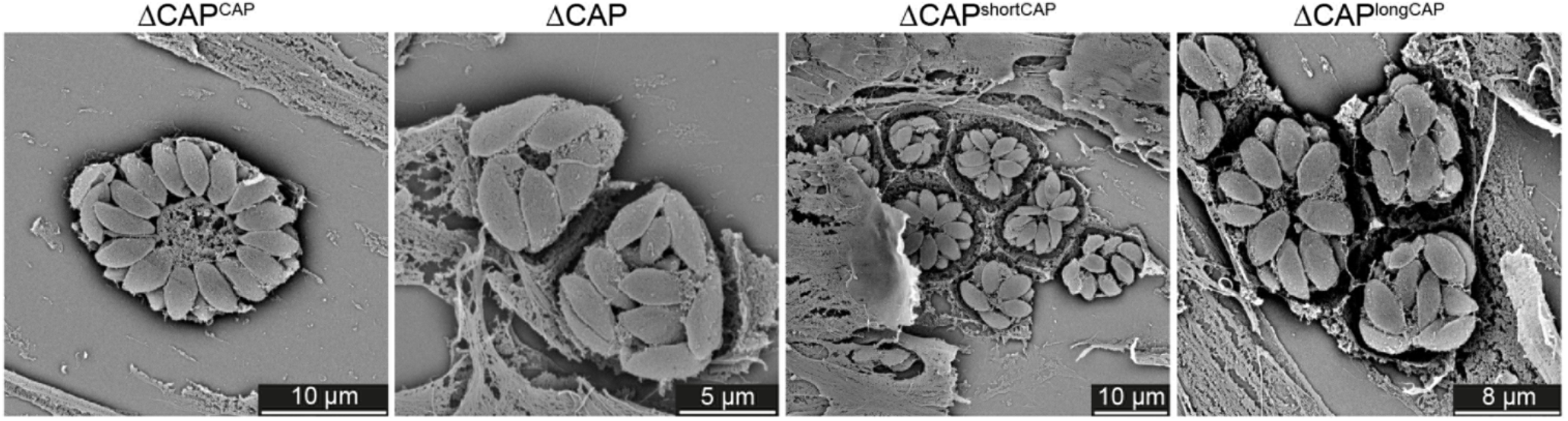
Scanning electron micrographs of intracellular ΔCAP and complemented lines. Vacuole organisation is disrupted in ΔCAP and ΔCAP^longCAP^ parasite strains.

**Supplementary Figure 8.**
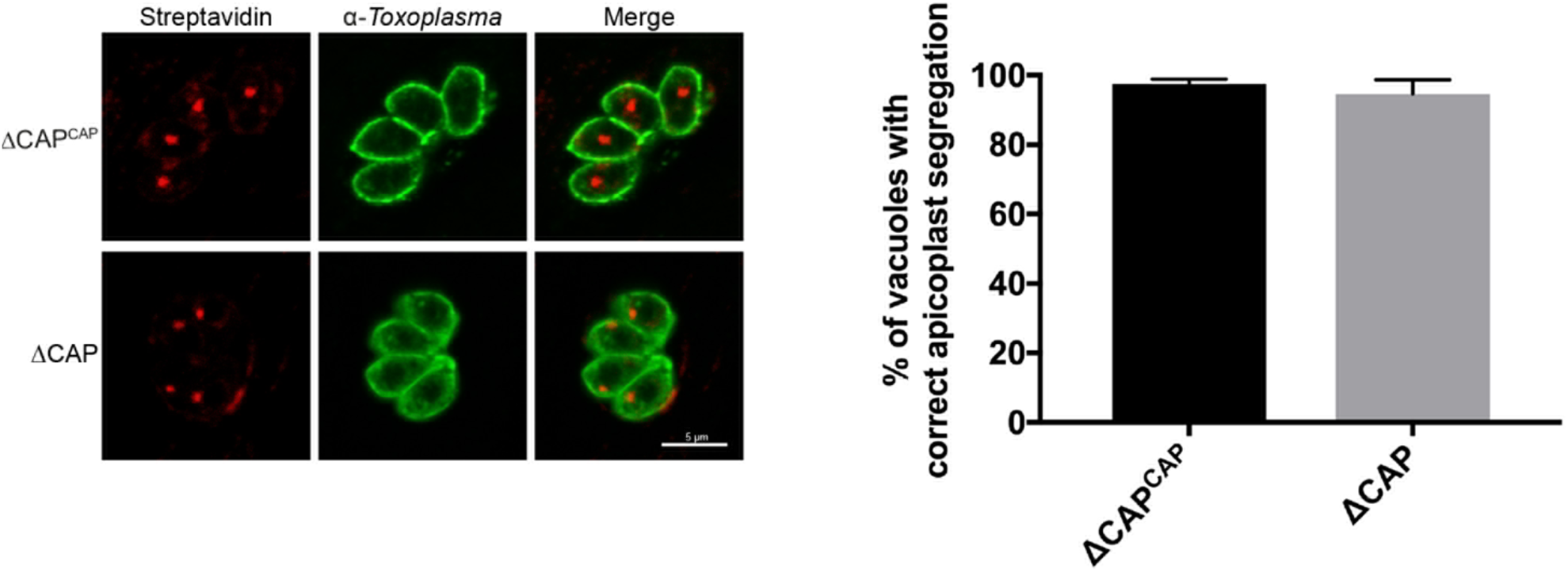
Apicoplast segregation analysis. Apicoplast segregation was not affected in the ΔCAP line. Representative IFA images (left) and quantification of apicoplast segregation (right). Data are represented as mean ± s.d. (*n*=3). Significance was assessed using an unpaired two-tailed *t*-test. Scale bar, 5 μm.

**Supplementary Figure 9.**
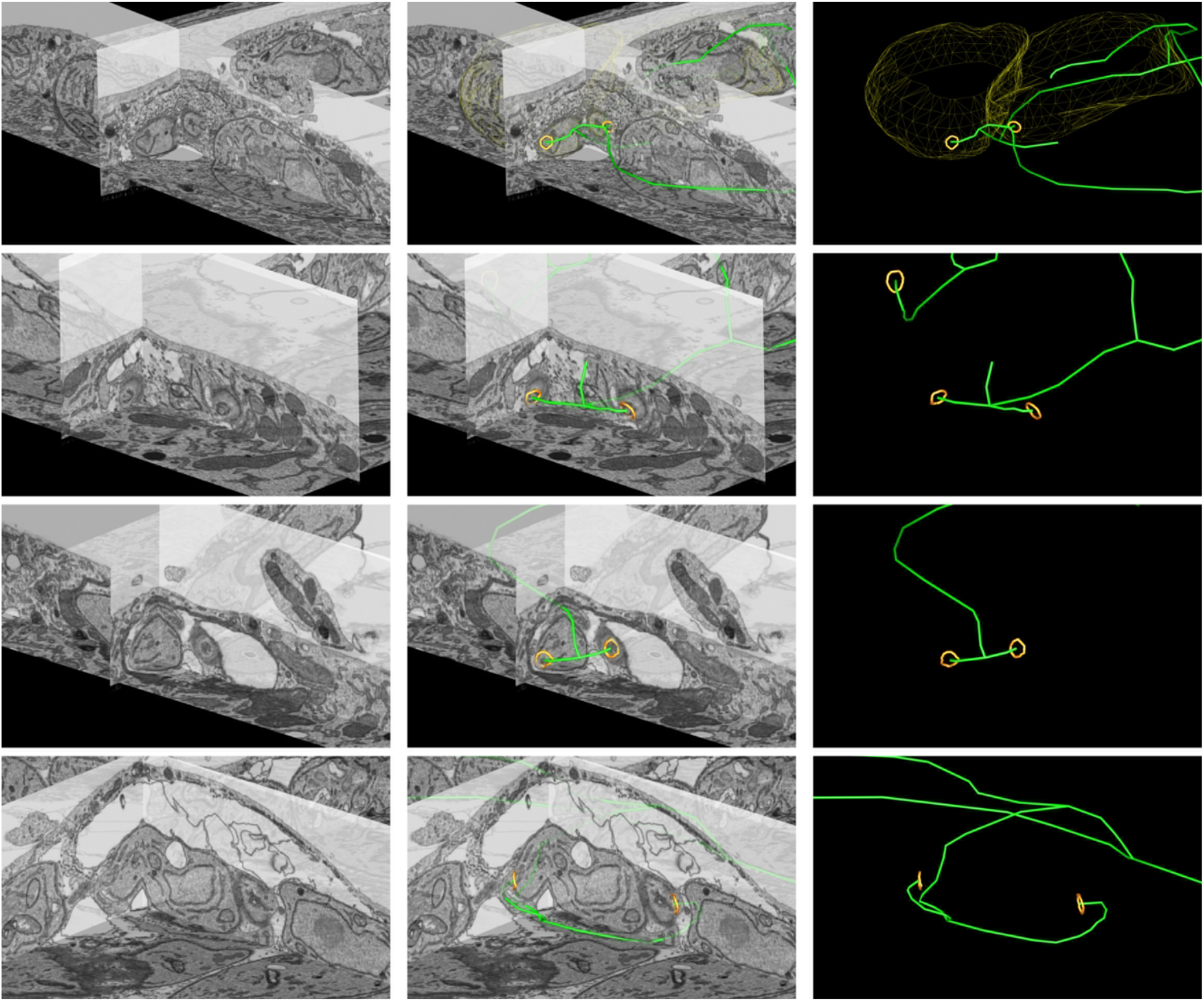
Close up FIB SEM images of parasite connections in the ΔCAP vacuole from (Fig. 7C-E) The 3D model highlights the residual body (green skeleton representing how the approximate centre can be traced through to the basal poles), parasite openings at the basal pole (orange ring) and coarse segmentation of two of the parasites (yellow). Scale bars are not shown since these are oblique views – see Fig. 7 and Methods for scale information.

**Video 1. Live egress imaging of a ΔCAP^CAP^ vacuole**

Egress was induced by addition of 50 μM BIPPO at 0 s. Image taken every 1.8 s. The video is played at 6.6 fps and the time is indicated in seconds. Scale bar, 10 μm.

**Video 2. Live egress imaging of a ΔCAP vacuole**

Egress was induced by addition of 50 μM BIPPO at 0 s. Image taken every 1.8 s. The video is played at 6.6 fps and the time is indicated in seconds. Scale bar, 10 μm.

**Video 3. Live egress imaging of a ΔCAP^shortCAP^ vacuole**

Egress was induced by addition of 50 μM BIPPO at 0 s. Image taken every 1.8 s. The video is played at 6.6 fps and the time is indicated in seconds. Scale bar, 10 μm.

**Video 4. Live egress imaging of a ΔCAP^longCAP^ vacuole**

Egress was induced by addition of 50 μM BIPPO at 0 s. Image taken every 1.8 s. The video is played at 6.6 fps and the time is indicated in seconds. Scale bar, 10 μm.

**Video 5. FRAP on a ΔCAP^CAP^ vacuole**

A single parasite was photobleached at 3 s. Images taken every 1 s. The video is played at 2 fps and the time is indicated in seconds. Scale bar, 5 μm.

**Video 6. FRAP on a ΔCAP vacuole**

A single parasite was photobleached at 3 s. Images taken every 1 s. The video is played at 2 fps and the time is indicated in seconds. Scale bar, 5 μm.

**Video 7. FRAP on a ΔGra2 vacuole**

A single parasite was photobleached at 3 s. Images taken every 1 s. The video is played at 2 fps and the time is indicated in seconds. Scale bar, 5 μm.

**Video 8. FIB SEM of the ΔCAP^CAP^ vacuole from Fig. 7A-B with 3D model overlay**

All FIB SEM slices of the ΔCAP^CAP^ vacuole Fig. 7A-B. The 3D model highlights the residual body (green skeleton representing how the approximate centre can be traced through to the basal poles, along with a structure that extends from that centre but has no connection) and parasite openings at the basal pole (orange ring). The volume of the FIB SEM dataset shown is indicated in the Materials and methods.

**Video 9. FIB SEM of the ΔCAP vacuole from Fig. 7C-E with 3D model overlays**

All FIB SEM slices of the ΔCAP vacuole Fig. 7C-E. The 3D model highlights the residual body (green skeleton drawn through the approximate central axis of the lumen of the connections between parasites), parasite openings at the basal pole (orange ring), coarse segmentation of the central part of two of the parasites (yellow), and part of the putative ER (blue, note that only a selected region was segmented; from the inner face of two of the basal pores through part of the decentralised residual body). The volume of the FIB SEM dataset shown is indicated in the Materials and methods.

**Supplementary File 1. Primers listed in the Materials and methods**

**Supplementary File 2. Synthetic DNA listed in the Materials and methods**

**Supplementary File 3. BioWave program details for FIB SEM**

